# NOTCH assembles a transcriptional repressive complex containing NuRD and PRC1 to repress genes involved in cell proliferation and differentiation

**DOI:** 10.1101/513549

**Authors:** Cecile M. Doyen, David Depierre, Ahmad Yatim, Alex Heurteau, Jean Daniel Lelievre, Yves Levy, Olivier Cuvier, Monsef Benkirane

## Abstract

NOTCH1 is best known as a master regulator of T-cell development with a strong oncogenic potential in developing T-cells. Upon induction of Notch, cells go through major transcriptional reprogramming that involves both activation and repression of gene expression. Although much is known about the transcriptional programs activated by Notch, the identity of the genes silenced downstream of Notch signaling and the mechanisms by which Notch down-regulates their expression remain unclear. Here, we show that upon induction of Notch signaling, ICN1-CSL-MAML1 ternary complex assembles a transcriptional Notch Repressive Complex (NRC) containing NuRD and PRC1. Genome wide analysis revealed set of genes bound and transcriptionally repressed by the NRC. Remarkably, among those genes, we found master regulators of cell differentiation and cell proliferation such as PAX5, master B-cell regulator and the DNA-binding transcriptional repressor MAD4. We propose that Notch possesses a dual role as direct activator and repressor by serving as a platform for the recruitment of co-activators and co-repressors on target genes and that both activities are required for Notch nuclear functions.

## INTRODUCTION

Notch signaling pathway is cell to cell communication mechanism that plays a key role in determining cell fates throughout embryonic development and in adult tissues. Consistently, dysfunctions in the Notch signaling pathway are associated with various human diseases including inherited genetic disorders and cancers. Human NOTCH1 was discovered in leukemic T-cells and was subsequently shown to be a master regulator of T-cell development with a strong oncogenic potential in developing T-cells (Aifantis et al., 2008; Weng et al., 2004). In addition to its oncogenic role in human T-cell acute lymphoblastic leukemia (T-ALL), Notch signaling drives the growth of a wide range of hematopoietic malignancies and solid tumors (South et al., 2012). Unexpectedly, Notch signaling can also act as tumor suppressor since mutations inactivating Notch pathway were reported in human cancer (South et al., 2012). Notch signaling has a remarkably simple and unusual signal-transduction framework. Its molecular architecture involves a small number of core signaling components, with no secondary messengers or signal amplification. Indeed, NOTCH protein is the transmembrane receptor, the cytoplasmic transducer and the nuclear effector of the pathway. The second unusual aspect is that Notch ligands are also transmembrane proteins, thus limiting signaling activation to direct cell-to-cell contacts. Activation of the Notch signaling pathway results in the sequential proteolysis of the NOTCH receptor by the ADAM protease and γ-secretase complex leading to release of the NOTCH Intracellular domain (Intracellular Notch - ICN) from the plasma membrane (Andersson et al., 2011; Bray, 2006; Guruharsha et al., 2012). ICN is then transported to the nucleus where it forms a ternary complex, with the DNA-binding protein CSL (RBP-Jκ/CBF-1) and the Mastermind family protein MAML1 (Andersson et al., 2011; Borggrefe and Liefke, 2012; Bray, 2006). Both CSL and MAML1 act as central components of Notch signalling by targeting nuclear ICN to Notch-responsive genes and by acting as coactivator, respectively. The assembly of the ICN-CSL-MAML ternary complex is required for most known Notch functions in both physiological and pathological contexts. In response to Notch signaling activation, cells go through major transcriptional reprogramming that involves both activation and repression of gene expression. Indeed, ICN-CSL-MAML1 serves as platform for the assembly of transcriptional activating complex containing several classes of transcriptional regulators including histone modifiers and RNA polymerase II recruiter (Bray et al., 2005; Collins et al., 2014; Fryer et al., 2004; Liefke et al., 2010; Saint Just Ribeiro et al., 2007; Yatim et al., 2012).

Although, ICN1 is most known as transcriptional activator, its translocation to the nucleus is also associated with gene transcriptional repression. For instance, ICN1 expression in hematopoietic progenitors suppresses myeloid specific genes (de Pooter et al., 2006; Kawamata et al., 2002), while its expression in B-lymphocytes represses several B-cell specific genes. Notch suppressive effect is believed to result from indirect mechanism, probably mediated by the expression of downstream transcriptional repressors, such as HES1 and HEY1. Here, we show that Notch signaling pathway regulates the assembly of a Notch-repressive complex (NRC) containing ICN1-CSL-MAML1, the Nucleosome Remodeling histone Deacetylase complex (NuRD) and the Polycomb Repressive Complex 1 (PRC1). The NRC is recruited to specific subset of genes to repress their transcription. Among the NRC repressed genes, we found important regulators of Notch function in T-cells.

## RESULTS

### Activation of Notch signaling pathway results in the assembly of complex containing CSL-ICN1-MAML1, the NuRD and the PRC1 subunits

We previously took a high-throughput proteomic inventory of the nuclear partners of the intracellular NOTCH1 (ICN1) (Yatim et al., 2012). ICN1 pulled down subunits from the NuRD (i.e.: HDAC1, MI2β, RBBP4, and GATAD2B), and the PRC1 (i.e.: RING1, RNF2). This study suggested that, in addition to the assembly of ICN1 transcriptional activating complex (NAC) (Yatim et al., 2012), ICN1 also interacts with transcriptional repressive complexes and could act as transcriptional repressor. We first sought to confirm the potential interactions between ICN1 and the subunits of the NuRD and PRC1 complexes identified by mass spectrometry. We performed immunoprecipitation (IP) with ICN1-FLAG and recovered the NuRD subunits HDAC1, MI2β/CHD4, MTA2 and MTA1, and the PRC1 subunits RING1 and BMI1 (Figure S1A) in addition to the well-known ICN1 interacting partners CSL and MAML1, FRYL and BRG1 as subunits of the NAC, fully confirming our earlier mass spectrometry screen (Yatim et al., 2012). Then we confirmed our results with a reverse IP in which we aimed to pull down ICN1 by using a FLAG tagged known member of repressive transcriptional complexes, HDAC1 (Figure 1A). The choice of HDAC1 was guided by its role as a subunit in different repressive complexes: NuRD, PRC2 and CSL-mediated repressive complex. Flag-HDAC1 IP successfully recovered its known interactors CSL, NuRD and PRC2 subunits. Interestingly, it also recovered ICN1, MAML1 and subunits of the PRC1 complex (Figure 1A). These results consolidate the existence of interactions between NuRD, PRC1, and NOTCH ternary complex.

**Figure 1.**
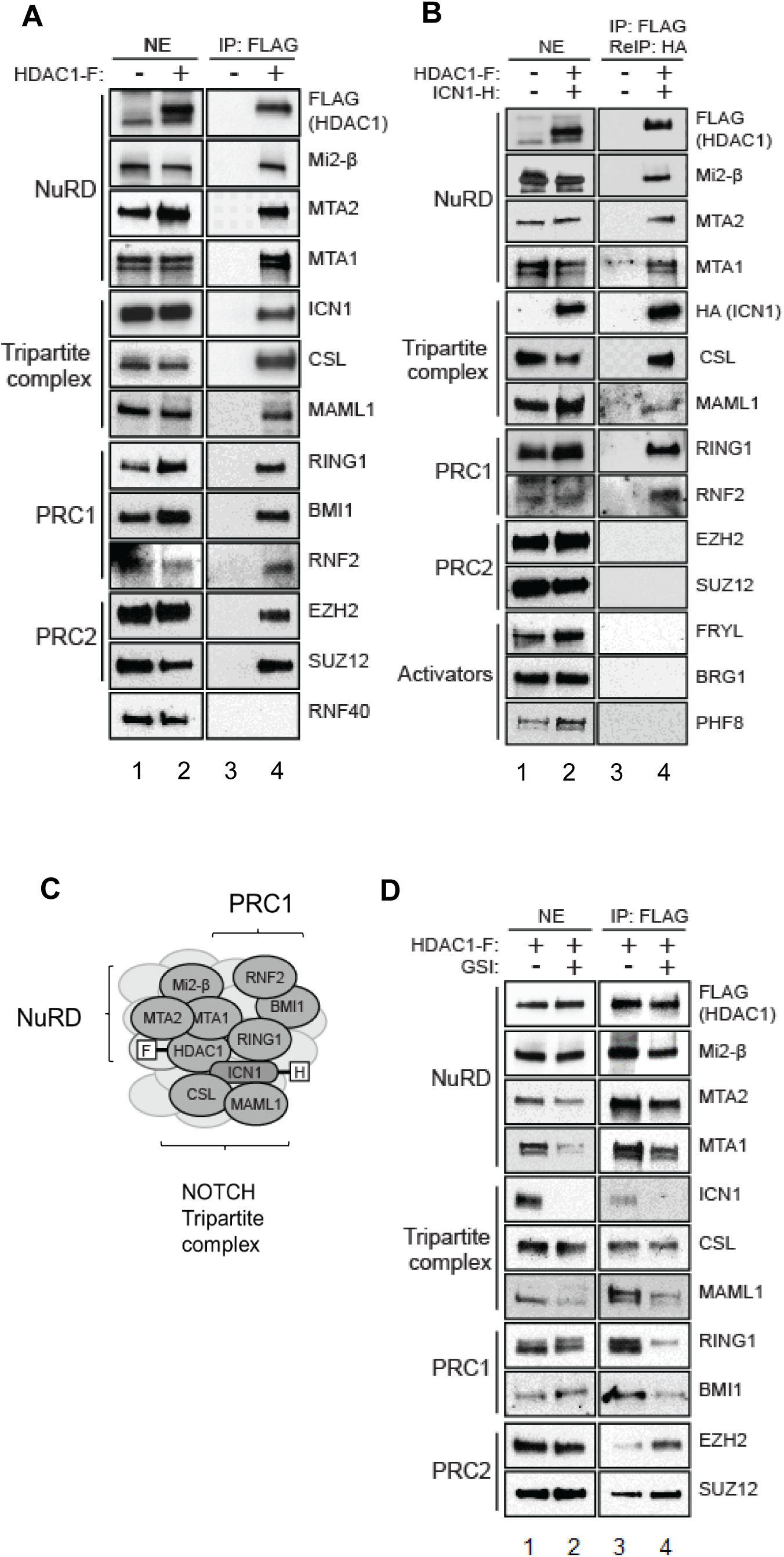
Activation of Notch signalling pathway induces the assembly of a transcriptional repressive complex containing CSL-ICN1-MAML1, NuRD and PRC1. **(A)** Immunoprecipitation of HDAC1-FLAG in SupT1 T-ALL cells, stably expressing FLAG-tagged HDAC1, analyzed by immunoblot. IP, immunoprecipitation; NE, nuclear extract. **(B)** Immunoprecipitation of HDAC1-FLAG followed by a second immunoprecipitation of ICN1-HA (ReIP) in SupT1 cells, stably expressing FLAG-HDAC1 and HA-tagged ICN1, analyzed by immunoblot. **(C)** A schematic model representing a complex containing Notch ternary complex, NuRD, and PRC1 subunits. **(D)** Immunoprecipitation of HDAC1-FLAG in SupT1 cells stably expressing FLAG-HDAC1. SupT1 cells were treated for 2 days with DMSO or GSI. See also Figure S1.

To test if the above-identified interactions reflect the presence of one or several complexes, we performed sequential IP experiments. SupT1 cells were engineered to express both Flag-HDAC1 and HA-ICN1. Flag and HA sequential IP using nuclear extracts was performed. As shown in figure 1B, subunits of the NuRD, PRC1 complexes together with CSL and MAML1 were found in Flag-HDAC1/HA-ICN1 IPs. None of the PRC2 subunits nor proteins known to associate with ICN1 activating complex (FRYL, BRG1 and PHF8) were recovered. This experiment shows that ICN1-CSL-MAML1 complex interacts with HDAC1 in either a large complex containing both NuRD and PRC1 or in two separate complexes. To verify this hypothesis, Flag-HDAC1 elutes were subjected to sequential IP using antibodies against the PRC1 subunit RING1. The presence of the NuRD subunits in IPed material was analyzed by western blotting (Figure S1B). Both MI2β/CHD4 and MTA2 were recovered. Moreover, endogenous RNF2 IP recovered the PRC1 subunits, NuRD subunits and ICN1-CSL-MAML1 (Figure S1C). Altogether, these experiments demonstrate that ICN1-CSL-MAML1, the NuRD and PRC1 assemble into a single complex (Figure 1C).

Having established the existence of a single complex containing ICN1-CSL-MAML1, NuRD and PRC1, we next sought to test the contribution of NOTCH signaling pathway to its assembly. For this, we inhibited NOTCH cleavage and nuclear translocation with the γ-secretase inhibitor GSI. GSI-treatment did not affect the levels of NuRD, PRC1, PRC2 subunits and CSL while it reduced nuclear accumulation of ICN1 and MAML1 (Figure 1D. compare lane 2 to 1). GSI-treatment did not affect the binding of the NuRD complex subunits to HDAC1 (Figure 1D compare lane 4 to 3). Remarkably, GSI reduced the binding of ICN1, MAML1 and PRC1 subunits, while it enhanced that of PRC2 subunits to FLAG-HDAC1 (Figure 1D compare lane 4 to 3). This experiment shows that activation of Notch induces the assembly of the large complex containing ICN1-CSL-MAML1, NuRD and PRC1 to the detriment of the NuRD-PRC2 complex. Taken together, our data show that upon activation of Notch signaling pathway ICN1-CSL-MAML1 ternary complex can assemble not only into a transcriptional activating complex (Yatim et al., 2012) but also into a transcriptional repressive complex containing NuRD and PRC1.

### Genome wide analysis defines the targets of the NOTCH transcriptional repressive complex

We sought to test if the interaction of NOTCH with NuRD and PRC1 might be detected in the context of their chromatin sites. Towards this goal, we performed chromatin IP coupled to high throughput sequencing (ChIP-seq) (Wang et al., 2008) for subunits of NuRD (MI2β, HDAC1, and MTA2,) PRC1 (BMI1 and RING1), CSL and ICN1, in T-ALL cell lines. We confirmed that the overlap between >20,000 ICN1 binding sites or ‘peaks’ and those identified for CSL was highly significant (Figure S2A; p-value < 1e-300). We also found a high overlap of binding sites between the NuRD complex subunits, MI2β, HDAC1, and MTA2) (Figure S2B; p-value < 1e-300). Finally, a lower fraction of shared binding sites was found between RING1 and Bmi1 (Figures S2C-D; p-value < 1e-198), as expected from the contribution of Bmi1 as Polycomb Group Ring Finger 4 (PCGF4), that assemble into PRC1.4, one of the PRC1 family complexes (Gao et al., 2012). Interestingly, only shared ICN1 and CSL-binding sites overlapped significantly (1,924 binding sites; p-value < 1e-5) with both NuRD and PRC1 binding sites (compare Figure S2E to S2F). We found that all subunits of the NRC: ICN1 and CSL, NuRD (HDAC1, MI2β, MTA2) and PRC1 (BMI1 and RING1) co-localized over 421 genomic sites, which represented a significant overlap (Figure 2A; p-value < 1e-33). In addition, GSI-mediated inactivation of the Notch pathway decreased such number of common NOTCH/NuRD/PRC1 sites by > 2.5 fold (Figure 2B; 164 sites), thereby impairing their significant overlap (p-value =0.99). BMI1 appeared to be the subunit whose peaks were most efficiently impaired upon GSI (Figure 2C; > 79 % of peak reduction), strengthening the view that Notch specifically recruits PRC1.4 complex. Supporting these results, BMI1, RING1 and NuRD subunits are specifically recruited to NOTCH bound sites and ICN1 and BMI1 binding were most specifically impaired upon GSI treatment (Figure 2D). Since PRC1 complexes recognize the repressive mark H3K27me3, we asked whether variations in H3K27me3 might account for variations in BMI1/PRC1 binding of NOTCH. We observed that H3K27me is not enriched at Notch-bound TSS. In addition, GSI did not affect H3K27me3 levels highlighting a role for ICN1, rather than H3K27me3, in PRC1 recruitment (Figure 2D). Taken together, our data show that the binding to chromatin of the NRC is dependent on NOTCH activation.

**Figure 2.**
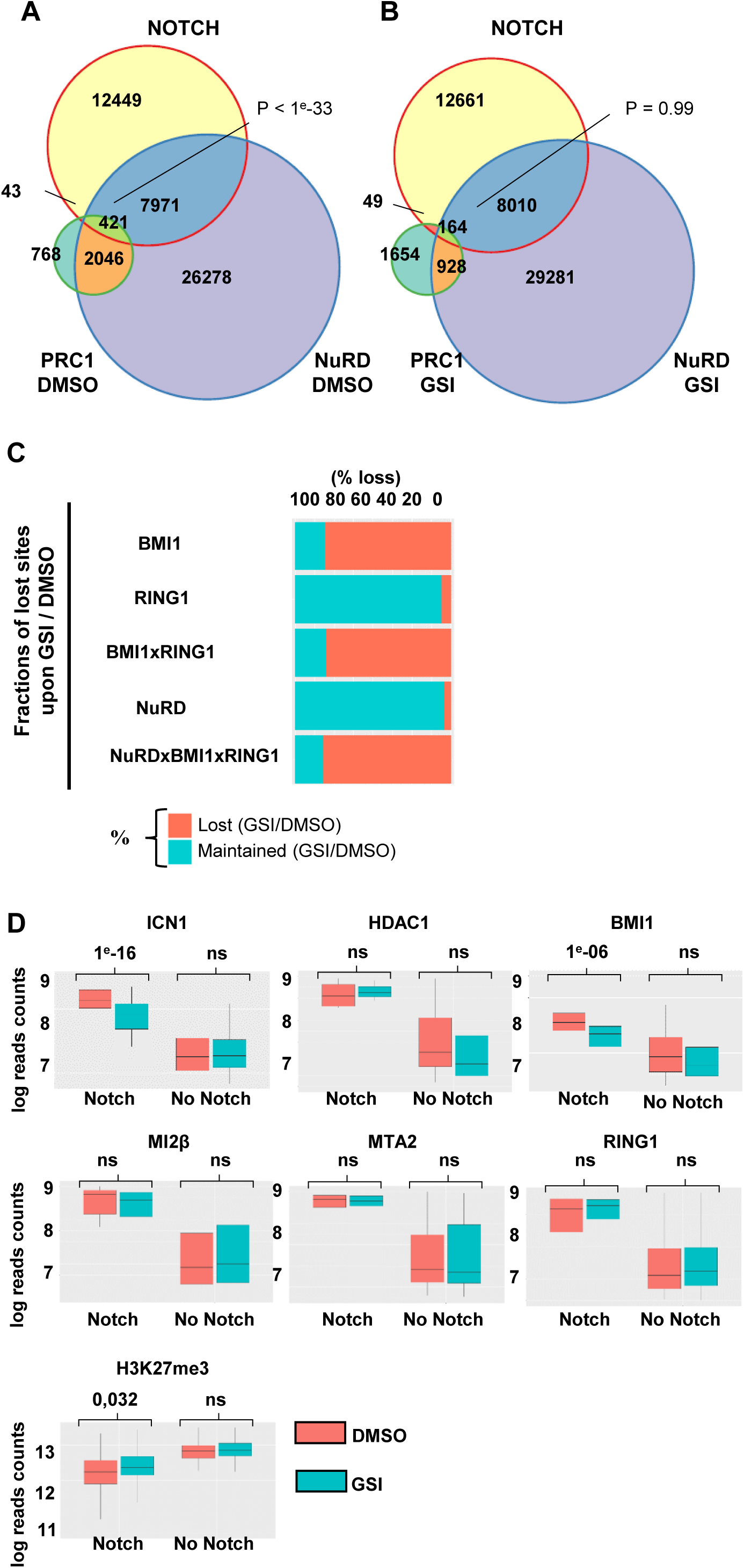
NOTCH co-localizes with NuRD and PRC1 at the promoters of thousands of genes. **(A-B)** Venn diagrams showing the significant overlap among genomic binding sites of NOTCH, NuRD and PRC1 in control cells (**A**) or in Gamma-secretase inhibitor (GSI) treated cells (**B**). ‘NOTCH’, ‘NuRD’ and ‘PRC1’ binding sites represent the overlapping sites where all Notch (CSL and ICN1), NuRD (HDAC1, MI2β, and MTA2) and PRC1 (RING1 and BMI1) subunits were co-localized (See Fig.S2B-D). P-value was calculated using Fisher exact test. **(C)** A graph representing the Notch-dependent loss of BMI1/RING1 and NuRD binding, as estimated by counting the proportion of peaks that are lost (red) or maintained (blue) upon GSI treatment compared to control (DMSO). **(D)** Box plots showing the variations in binding levels (normalized ChIP-seq reads) of the indicated factors upon treatment with GSI (blue) as compared to DMSO (red). Y-axis: Levels of normalized ChIP-seq reads over input for the indicated factor. See also Figure S2.

### The presence of the NRC on chromatin is associated with transcriptional repression of the target genes

The above data demonstrate that activation of the Notch pathway recruits the NRC to chromatin, which raises the possibility that NOTCH may act as a direct transcriptional repressor of the targeted genes. To test this hypothesis, we performed a combinatory analysis of RNA-seq and ChIP-seq data obtained from T-ALL cells that were mock- or GSI-treated. This approach revealed several gene categories among NOTCH target genes. Of the 5051 genes bound by ICN1 at +/-1kb from TSSs, 1745 were differentially expressed (DE) upon inhibition of Notch pathway by GSI compared to control (Figure 3A). The genes were either activated (2062) or repressed (2058) upon NOTCH inactivation by GSI. ICN1 binding alone could not predict activation or repression (odds ratio of 1), whereas NRC binding specifically increased the odds ratio of Notch-repressed genes by 3.7 fold over Notch-activated genes (Figure S3B). As such, Notch-mediated repression was encountered when genes were bound by NRC subunits BMI1, RING1 and NuRD (Figures 3B, p-value < 1e-4; Figure S3A-B). In stark contrast, binding of ICN1 in absence of NuRD and PRC1 was associated with transcriptional activation (Figure 3B, p-value < 1e-4). This experiment establishes that direct transcriptional repression by Notch occurs in T-cells and represents more than 50% of Notch transcriptional responses.

**Figure 3.**
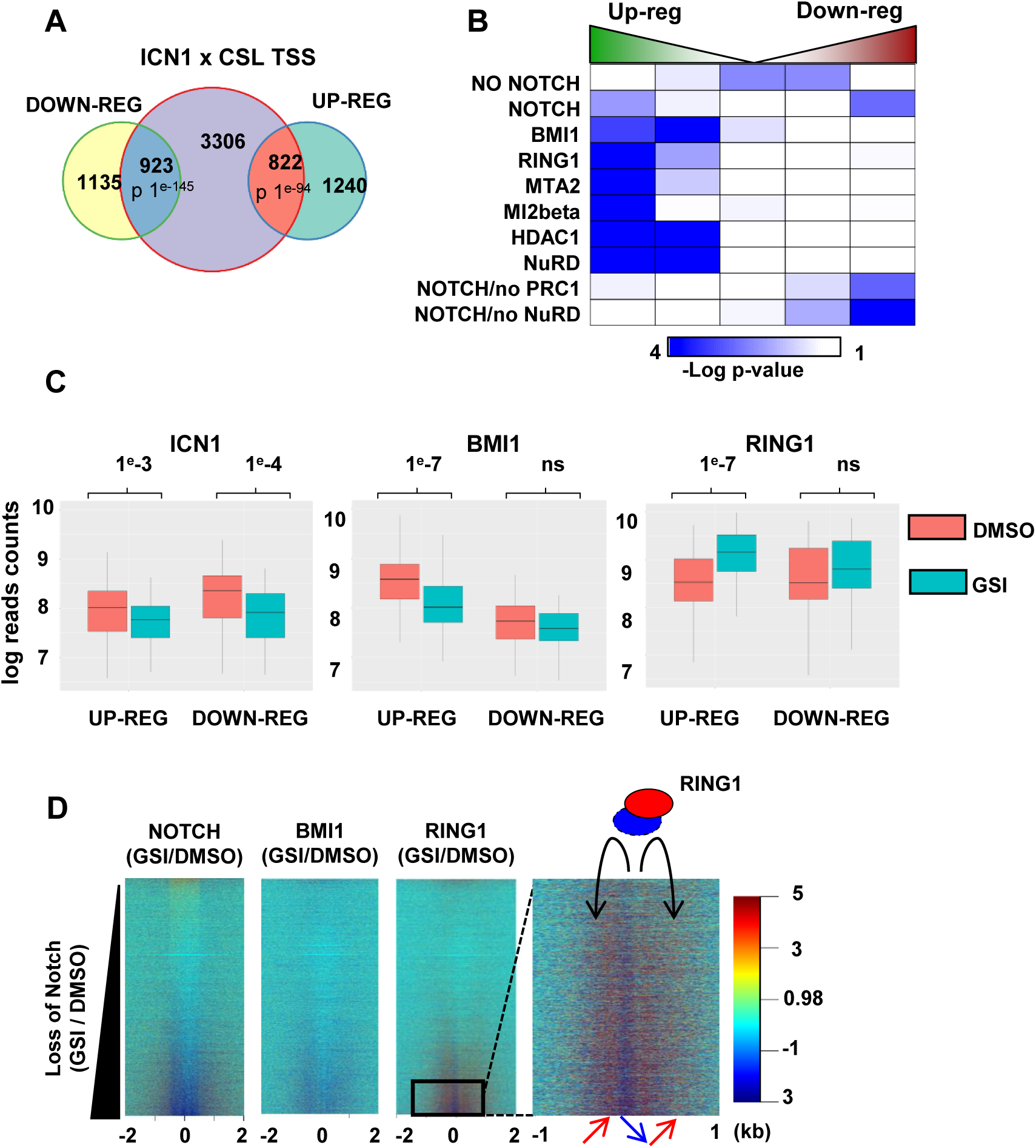
The NRC is a transcriptional repressor of specific subset of genes. **(A)** A Venn diagram showing intersection between lists of genes harboring the binding sites of both ICN1 and CSL on their TSSs and lists of genes up- or down-regulated as obtained by RNAseq in GSI-vs DMSO-treated cells. P-value was calculated using Fisher exact test. **(B)** Intersection matrix showing the statistical enrichment of NOTCH-bound genes and the co-localization with NuRD, and PRC1 subunits, depending on the gene expression upon GSI treatment. **(C)** Box plots showing the variations in binding levels (normalized ChIP-seq reads) of ICN1, BMI1, RING1 subunits for genes that are up-regulated or down-regulated upon GSI (blue) as compared to DMSO (red). **(D)** Heatmaps showing the net difference in normalized ChIP-seq reads in cells treated with GSI compared to DMSO, for NOTCH, BMI1, and RING1. The net variations in reads were aligned over TSS to estimate dynamics depending on the gene ranking from increasing to decreasing NOTCH binding level at the TSS (+/-1 kbp). See also Figure S3.

Next, we specifically analyzed the dynamics of the co-factors binding upon Notch inactivation by GSI within the two sets of NOTCH-target genes (Figure 3C). As expected, ICN1 recruitment was impaired upon GSI treatment for both repressed and activated genes (Figure 3C). Bmi1 is specifically enriched on Notch-repressed genes. Like Bmi1, RING1 dynamics was encountered specifically for up-regulated genes (Figure 3C). Of note, the levels of RING1 increased only if Notch co-localized with Bmi1 (Figure 3C) thus showing that Notch-mediated regulation of PRC1 binding was specific to PRC1 complex containing BMI1. Further measures of GSI-mediated changes in ICN1 binding and in gene de-regulation show that 62/86 up-regulated genes, associated with BMI1 binding, were lost upon GSI treatment (Figures S3C). Thus, NOTCH-mediated transcriptional repression involves a dynamic behavior of PRC1. Together, our data show that upon activation of NOTCH signaling, the NRC binds to and silences NOTCH target genes.

To understand the Notch-dependent dynamics of RING1, we ranked genes according to the effect of Notch inhibition by GSI and analyzed the binding of ICN1, BMI1 and RING1 (Figure 3D). Consistently, the decrease in Notch binding upon GSI treatment was accompanied by a large decrease in Bmi1 binding and concomitant redistribution of RING1 from TSS to surrounding chromatin +/-1kb (Figure 3D, arrows). Therefore, impairing Notch binding induces a dynamic re-localization of PRC1 that accompanies a severe reduction of BMI1 binding at TSSs (Figure S3D). Of note, Notch-dependent dynamic of RING1 and BMI1 binding, in which the RING1 redistribution mirrors the loss of BMI1, occurs specifically at genes repressed by a distant NOTCH bound enhancer (Figure S3E). Taken together, our data highlight NOTCH signaling-dependent dynamics of the PRC1 binding at NOTCH repressed genes.

### Notch-bound enhancers regulate distant genes through long-range contacts

ICN1 binding was also detected at enhancers (Figure S4A). We thus sought to assess the respective influence of NOTCH and co-factors depending on their association with enhancer regions. NOTCH, NuRD, and PRC1 significantly overlap at enhancer sites (Figures S4A-C; p-value of 10e-216). Moreover, of the 1,464 NOTCH binding sites present at enhancers, only 16% did not co-localize with PRC1 or NuRD (Figure S4D), thereby raising the possibility that NuRD and PRC1 might participate in a NOTCH-mediated regulation of enhancer activity.

To test this hypothesis, we integrated promoter capture Hi-C data, a method to study long-range interactions in a 3D architecture of genome (Javierre et al., 2016). This identified all NOTCH-bound and NOTCH-unbound enhancers that established long-range contacts with their candidate target promoters, as illustrated for the NOTCH target gene PTPRC (Figure 4A). In absence of BMI1, promoters in direct contact with Notch-bound enhancers were not enriched for de-regulated genes upon ICN1 depletion (Figure 4B). In contrast, the association of BMI-bound promoters with NOTCH-bound enhancers had a clear negative impact on the contacted genes, as confirmed by the specific enrichment in up-regulated genes (Figure 4B; 6-7^th^ rows). Interestingly, such influence was detected even in absence of NOTCH binding at promoters (Figure 4B; 5^th^ row) showing that the influence of the NOTCH/PRC1/NuRD-bound enhancers depends on long-range contacts.

**Figure 4.**
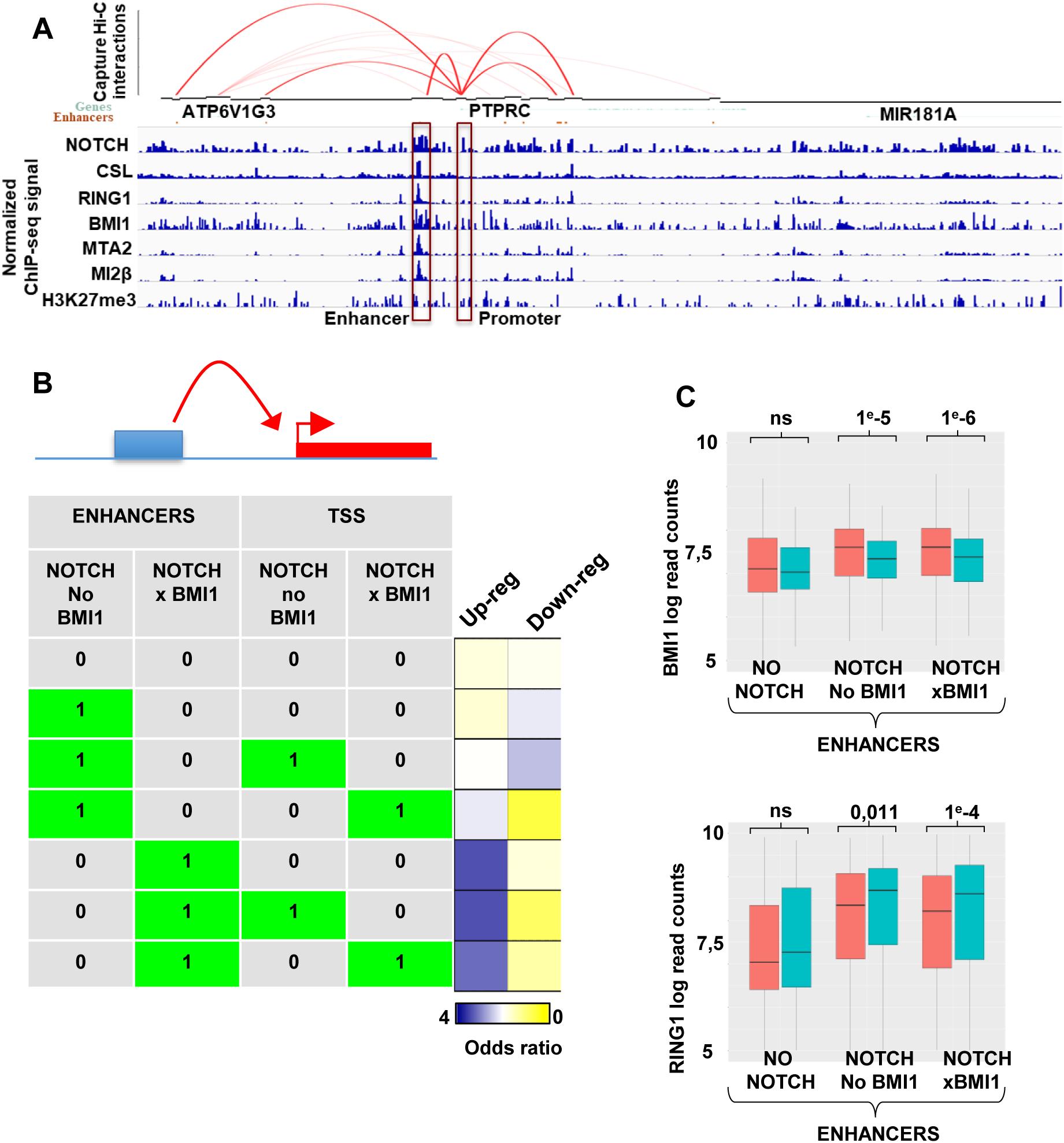
Long-range contacts between NOTCH-bound enhancers and promoters of Notch-target genes establish transcriptional repression. **(A)** Gbrowse view of promoter capture Hi-C data representing the long-range contacts between NOTCH-bound enhancers and the TSS of the NOTCH repressed gene PTPRC. ChIPseq profiles are shown for NOTCH, CSL, NuRD and PRC1 subunits and H3K27me3. **(B)** Scheme representing the enrichment tests of the Notch-bound enhancers or promoters and the NOTCH-BMI1-bound enhancers or promoters within the lists of up-and down-regulated genes obtained by DEseq2 after RNAseq from GSI-compared to DMSO-treated cells. Notch/Bmi1-bound enhancers were identified as enhancers that physically associate with TSSs through 3D chromatin loops, as detected by promoter capture Hi-C (see panel A for an example). **(C)** Box plots showing the dynamic changes in TSS binding of BMI1 and RING1 in GSI (blue) compared to DMSO (red), depending on 3D contacts with enhancers. Y-axis: Levels of normalized ChIP-seq reads over input. See also Figure S4.

Next we analyzed the dynamics of the BMI1 and RING1 binding to the TSSs within promoters depending on contacts with Notch-bound enhancers or not. Loss of ICN1 binding to its target enhancers upon GSI-treatment was accompanied by the loss of BMI1 binding to TSSs within promoters contacted by ICN1-bound enhancers, regardless of the presence of BMI1 at the enhancers (Figure 4C, 1e-5). RING1 binding increased upon GSI-treatment in a BMI1/ NOTCH dependent manner (Figure 4C). No influence of GSI treatment on BMI1 and RING1 binding was observed when ICN1-unbound enhancers were considered (Figure 4C). Overall, our data establish that NOTCH-bound enhancers can regulate BMI1 and RING1 binding over the distant, physically associated TSS, thereby controlling gene expression.

### The Notch repressive complex regulates transcription of genes involved in cell differentiation and proliferation

The above biochemical and genome-wide analyses demonstrate that the ICN1-CSL-MAML1 tripartite complex assembles a repressive complex containing NuRD and PRC1 to repress a subset of genes in response to NOTCH signaling. Among the NOTCH-target genes that are up-regulated upon GSI, we selected 14 candidates for their role in cell development and proliferation. We performed quantitative RT-PCR (RT-qPCR) on total RNA prepared from mock- and GSI-treated T-ALL cells and revealed increased expression of the 14 candidate genes (Figure S5A). As a control, GSI-treatment reduced mRNA levels of the well-known NOTCH -activated genes HES, DTX and IL7R. To ensure that the observed effect is transcriptional, we performed 4sU RNA incorporation to label nascent transcripts before RTqPCR (Schwalb et al., 2016). As shown in Figure S5B, while GSI repressed transcription of HES, DTX, and IL7R, it enhanced that of the 14 candidate genes. Importantly, both CSL knockdown and overexpression of a MAML1 dominant-negative showed a similar effect to that observed using GSI, confirming that the transcriptional repression of these genes is dependent on the ternary complex (Figure S5C-E).

We next sought how the Notch-mediated transcriptional silencing may contribute to the Notch biological functions, namely cell differentiation and proliferation. We thus focused on two Notch target genes: the B-cell master regulator PAX5 and the MYC repressor MAD4. PAX5 is a B-cell specific transcription factor required for commitment of hematopoietic progenitors to the B-cell lineage (Medvedovic et al., 2011; Nutt et al., 1999; Urbanek et al., 1994). We hypothesized that NOTCH-mediated recruitment of co-repressors to the PAX5 locus might block its transcriptional activation and suppress cell commitment to the B-lineage. To test this, we quantified PAX5 mRNA and protein levels in three T-ALL cell lines (DND41, HPB-ALL, and SupT1) that were mock- or GSI-treated. While GSI treatment down-regulated the Notch-activated gene, *DTX* mRNA, it enhanced *PAX5* mRNA and protein in all the tested T-ALL cell lines (Figures 5A and Figure S5F). Using nascent transcript analysis, we confirmed a transcriptional de-repression of PAX5 upon GSI treatment (Figure 5B). Enhanced expression of PAX5 upon GSI treatment was accompanied by the activation of its target genes *RAG1, RAG2* and the pre-B Cell Receptor (pre-BCR) subunits *CD79a* and *VpreB* (Figure 5D) (Busslinger, 2004; McManus et al., 2011; Pridans et al., 2008). Interestingly, overexpression of ICN1 in the B-cell acute leukemia REH (B-ALL) not only repressed *PAX5* and its target genes *RAG1, RAG2* and *CD79a*, it also enhanced the expression of *TCF7*, a transcription factor critical for T-cell commitment and *DTX* (Figure 5E-F). To further assess PAX5 repression by NRC, we used CRISPR CAS technology to invalidate the CSL binding site within the PAX5 locus and re-evaluated its impact on NOTCH-mediated repression of PAX5. We obtained a clone CRIPSR57 that was specifically depleted from the CSL binding site (Figure S5I). Remarkably, the deletion of the CSL binding site is accompanied by loss of NOTCH-mediated transcriptional repression of PAX5 without affecting that of other Notch activated genes such as HES1 and DTX (Figure 5F). Therefore, our results demonstrate a specific role for the intragenic CSL binding site in Notch-mediated repression of Pax5, further illustrating our genome-wide analyses highlighting the repression of TSSs by distant enhancer-bound NRCs.

**Figure 5.**
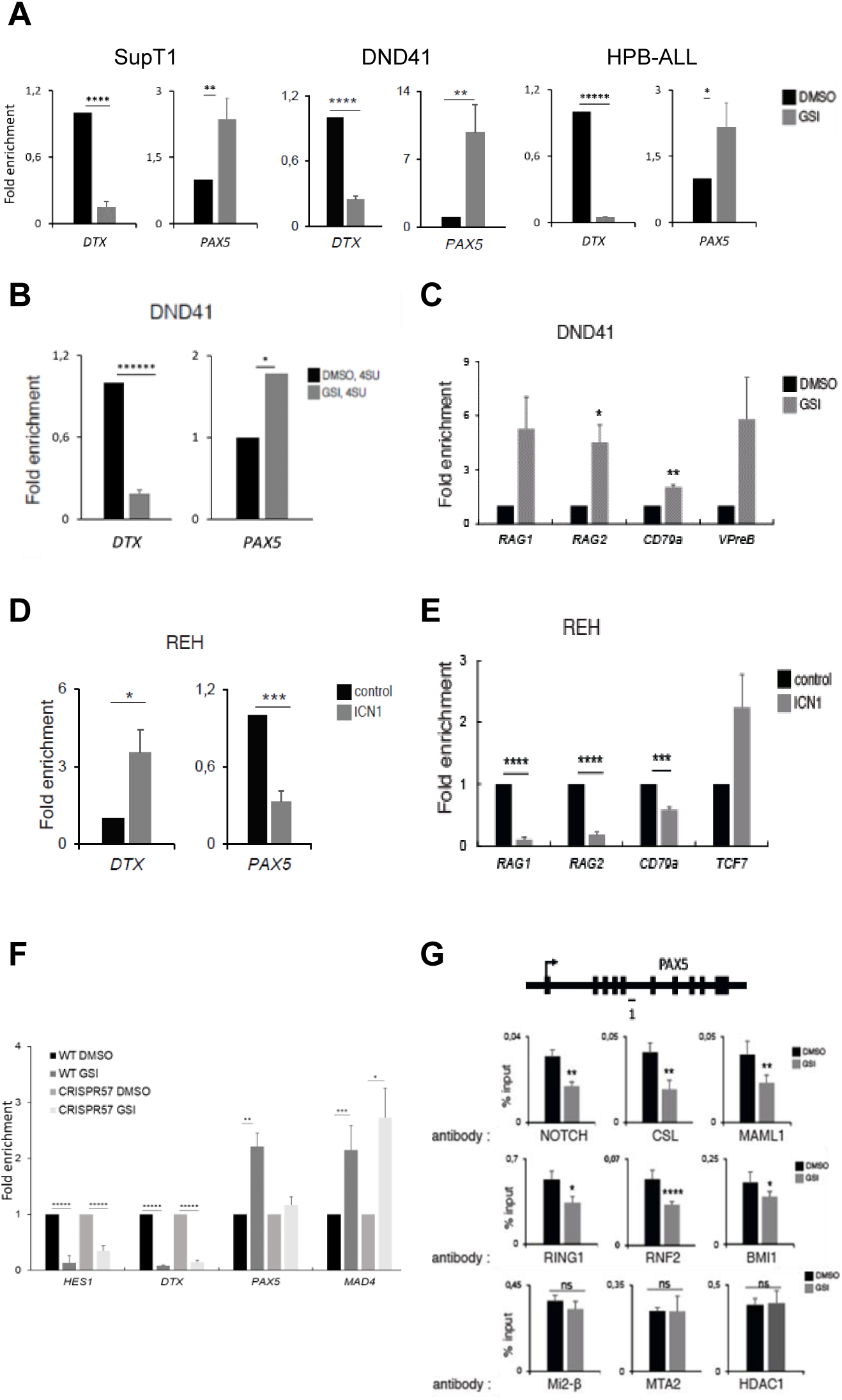
NRC is recruited to PAX5 and mediates its transcriptional repression. **(A)** *DTX* and *PAX5* measured by qRT-PCR in SupT1, DND41, and HPB-ALL cells treated 2 days with DMSO or GSI. **(B)** *DTX* and *PAX5* measurement by qRT-PCR in DND41 treated 2 days with DMSO or GSI and for15 min with 4sU. **(C)** qPCR measurement of PAX5-target genes mRNA levels in DND41 T-ALL cells. **(D)** Analysis of *DTX* and *PAX5* mRNA levels by qRT-PCR genes in REH B-ALL overexpressing control or ICN1. **(E)** *PAX5*-target genes and *TCF7* measured by qRT-PCR in REH B-ALL cells overexpressing control or ICN. **(F)** qPCR measurement of *HES1, DTX, PAX5* and *MAD4* mRNA in SupT1 wild-type and CRISPR57 cells treated 2 days with DMSO or GSI. **(G)** Analysis of locus occupancy of NOTCH, CSL and MAML1 (upper panel), PRC1 subunits (RING1; RNF2; BMI1) (middle panel) and NuRD subunits (MI2β; MTA2; HDAC1) (lower panel) on PAX5 gene by qChIP assay after treatment of SupT1 cells for 2 days with DMSO or GSI and the antibodies directed against. The CSL binding regions identified in *PAX5* locus were PCR amplified from the precipitated and input DNA. The position of the PCR amplicons is illustrated in the scheme (black dashes). The results are expressed as a percentage relative to input. mRNA levels were normalized to those of GAPDH (A-F). See also Figure S5.

Next, we asked whether the NOTCH transcriptional repressive complex is recruited to PAX5 locus. For this purpose, we performed ChIP-qPCR experiment in SupT1 cells that were mock- or GSI-treated and assessed for the recruitment of ICN1, CSL, MAML1 and subunits of the NuRD and PRC1. We found that ICN1, CSL, MAML1, NuRD and PRC1 subunits associated specifically with the same region of the *PAX5* locus containing a CSL binding site (Figure S5I). As previously described (Yatim et al., 2012), ICN1, CSL and MAML1 were present at the CSL binding site within HES1 and NOTCH3 (Figure S5H). GSI-treatment of SupT1 cells results in reduced binding of the tripartite complex to the *PAX5* gene (Figure 5H), and to HES and NOTCH3 (Figure S5J). Consistent with ChIP-seq data, inhibition of NOTCH signaling pathway by GSI results in reduced recruitment of the subunits of PRC1 to the *PAX5* locus (Figure 5H, middle panel). GSI, however, did not affect the levels of the NuRD subunits associated with *PAX5* (Figure 5H, lower panel). Taken together, these experiments show that upon Notch activation, ICN1-CSL-MAML1/NURD/PRC1 complex is recruited to *PAX5* locus via the CSL binding site to directly repress its transcription.

A second example of Notch repressed gene is the DNA-binding transcription repressor MAD4, a member of the MYC-MAX-MAD network. Whereas the MYC-MAX heterodimer induces activation of Myc-target genes and promotes cell proliferation, the MAX-MAD heterodimer acts as a repressor of MYC-MAX by competing for the same E-box sequence (Ayer and Eisenman, 1993; Grandori et al., 2000; Grinberg et al., 2004; Packard and Cambier, 2013). Given the importance of MYC in NOTCH-mediated cell proliferation (Palomero et al., 2006; Sanchez-Martin and Ferrando, 2017; Wang et al., 2011), we hypothesized that NOTCH might enhance MYC expression by direct transcriptional repression of MAD4. First, MAD4 expression was analyzed in three T-ALL cell lines both at the mRNA and protein levels. Enhanced MAD4 mRNA (Figure 6A and 6B) and protein (Figure 6C) levels was observed upon GSI-treatment of the T-ALL cells. Enhanced expression of MAD4 upon GSI treatment was accompanied by reduced expression of MYC and enhanced expression of MAX (Figure 6E). Taken altogether, our data therefore show that NOTCH pathway may impede on the MYC-MAX-MAD balance by favoring MYC expression through transcriptional silencing of MAD4.

**Figure 6.**
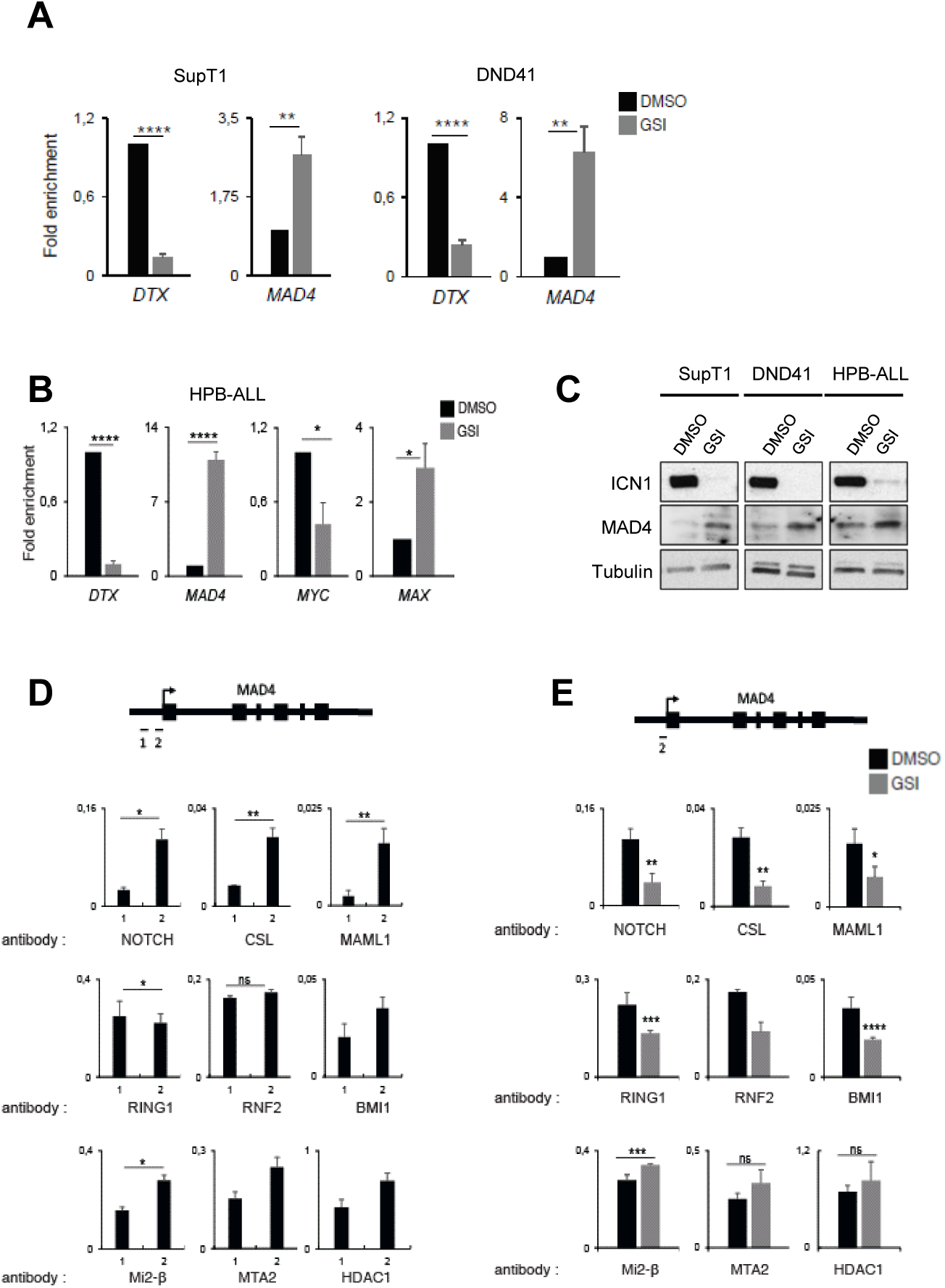
NRC is recruited to MAD4 promoter and mediates its transcriptional repression. **(A)** *DTX* and *MAD4* measurement by qRT-PCR in SupT1 and DND41 cells treated 2 days with DMSO or GSI. **(B)** Analysis of *DTX, MAD4, MYC* and *MAX* mRNA levels by qRT-PCR in HPB-ALL cells treated 2 days with DMSO or GSI **(C)** Immunoblot analysis of SupT1, DND41, HPB-ALL cells. Cells were treated 5 days with DMSO or GSI. **(D)** Analysis of locus occupancy of NOTCH, CSL and MAML1 (upper panel), PRC1 subunits (RING1; RNF2; BMI1) (middle panel) and NuRD subunits (MI2β; MTA2; HDAC1) (lower panel) on MAD4 gene by qChIP assay of SupT1 T-ALL cells. The positions of the PCR amplicons are illustrated by black dashes. **(E)** Same as (D). Cells were treated 2 days with DMSO or GSI.

We next analyzed the recruitment of the ICN1-CSL-MAML1, subunits of the NuRD and PRC1 to *MAD4* locus by a ChIP qPCR experiment. We designed two amplicons, only one of which contains the CSL binding site (Figure 6D). Consistently with our previous finding showing that the tripartite complex is required for the MAD4 repression, we confirmed that ICN1, CSL and MAML1 are bound onto the CSL binding site (Figure 6D, upper panel). Moreover, we also detected the binding of NuRD and PRC1 subunits to the same site, confirming the binding of NRC to MAD4 (Figure 6D, lower panels). Consistent with the data obtained by ChIP-seq and ChIP of PAX5, inhibition of the NOTCH signaling pathway leads to reduced recruitment of NOTCH tripartite complex and PRC1 subunits, but not that of NuRD subunits, at MAD4 locus (Figure 6E).

Altogether, our results show that upon activation of NOTCH signaling, the ICN1-CSL-MAML can serve as a platform for the assembly and recruitment of a transcriptional repressive complex to silence a subset of downstream target genes including those involved in cell proliferation and differentiation.

## DISCUSSION

The Notch signaling pathway is an evolutionarily conserved cell communication mechanisms which plays a prominent role in dictating cell-fate decision. Dysfunctions of the pathway are associated with various human diseases including inherited genetic disorders and cancers. In this study, we reveal that the CSL-ICN1-MAML ternary complex assembles into a transcriptional repressive complex containing, NuRD and PRC1. Such Notch-repressive complex (NRC) targets a specific subset of genes to repress their expression. Among the targeted genes, we found the B-cell master regulator PAX5 and the MYC repressor MAD4, highlighting the importance of the direct transcriptional repression by Notch in its biological outcome, namely cell differentiation and proliferation.

### Assembly and mechanism of transcriptional repression by the NRC

Three major findings emerge from our biochemical data. First, like the Notch activating complex (NAC), the assembly of the NRC is dependent on the activation of Notch signaling pathway. Indeed, inactivation of the Notch signaling resulted in the disassembly of the NRC. Second, although they achieve opposed function in gene regulation, both NAC and NRC are assembled around the ICN1-CSL-MAML1 ternary complex suggesting, yet to be identified, a regulatory mechanism orchestrating the formation of the two complexes. Recent study reported a physical and functional interaction between ICN1 and PRC2 using mouse MEF cells (Han et al., 2017). The fact that ICN1 interaction with the PRC2 was not recovered in T-ALL cell suggest a cellular context dependency. Third, the identification of the NRC revealed a previously undescribed interaction between NuRD and PRC1. We observed that GSI-mediated inactivation of ICN1 release results in loss of the interaction between PRC1 and NuRD, which was accompanied by increased interaction between the NuRD and PRC2 (Figure 1D). These experiments suggest that ICN1-mediated interaction between NuRD and PRC1 is at the expense of the interaction between NuRD and PRC2. Further biochemical analyses are required to decipher the molecular mechanisms regulating the inter-dependent dynamic of these chromatin repressive complexes and its impact on gene regulation.

Combining RNA-seq and ChIP-seq data we found that NRC is associated with transcriptionally repressed genes but not with the Notch activated genes. As expected, ICN1 and CSL were associated with both activated and repressed genes. Such data strengthen the conclusion that activation of the Notch signaling results in the concomitant assembly of both NAC and NRC that are subsequently targeted to their specific genes. Inactivation of Notch signaling results in the loss of ICN1 and PRC1 from their target genes leading to their transcriptional de-repression. Interestingly, we found that the presence of NuRD at NRC-repressed genes was unchanged upon inhibition of Notch signaling. Whether it reflects a partial dissociation of the NRC or a recruitment of an activating complex containing NuRD is to be uncovered. Indeed, it has been shown that the NuRD complex can function both as repressor and activator of transcription (Bornelov et al., 2018; Dege and Hagman, 2014; Hutchins et al., 2002; Williams et al., 2004; Yang et al., 2012). NRC-mediated transcriptional repression of its target genes involves both its direct recruitment to promoter region or to enhancers contacting promoter of targeted genes. We found a tight regulation of PRC1 binding upon Notch activation and an inter-dependence of RING1 and BMI1 binding to CSL binding sites. This tight regulation occurs both at promoters and enhancers of Notch-repressed genes. Remarkably, NRC binding to enhancers appears to be sufficient to inhibit the contacted promoter even if the latter is not bound by ICN1.

### Insights into the molecular mechanisms directing the ternary complex towards NRC or NAC assembly

Activation of the Notch signaling pathway results in the assembly of both NAC and NRC. Both of which are assembled around CSL-ICN-MAML1 ternary complex and recruited to their target chromatin through the CSL DNA binding motif. These results raise two important questions. First, what directs the ternary complex towards the assembly of the NAC versus the NRC? Second, what specify the target genes for both complexes? Like for other transcription factors (Freund et al., 2017; Huang et al., 2009; Van Nguyen et al., 2012), post-translational modifications (PTMs) of the ternary complex subunits may play a role. Indeed, PTMs of both ICN1 and MAML1 have been shown to regulate their transcriptional activity by modulating their stability and/or their network of interactions with co-activators or co-repressors (Andersson et al., 2011; Antila et al., 2018; Fortini, 2009; Lindberg et al., 2010; Popko-Scibor et al., 2011). Thus, analyses of PTMs of the ternary complex subunits associated with NAC and NRC together with the identification and the characterization of protein-modifying enzymes associated with both complexes will be of importance.

Consistent with previous studies, our ChIP-seq data indicates that thousands of genes are bound by ICN1, but only a small subset are differentially expressed in response to Notch activation. As specific combinations of transcription factors (Refs) often control the expression of tissues-specific genes, we hypothesized that combinatorial interaction between Notch and other transcription factors might determine the specificity of Notch transcriptional programs. Consistently, nuclear interactome of ICN1 revealed the presence of other transcription factors and lineage-specifying transcription factors (such as BCL11B, HEB, RUNX1 and IKAROS) (Yatim et al., 2012). Among them, the transcriptional repressor BCL11B is known to play a major role in T-cell commitment (Li et al., 2010a; Li et al., 2010b). It has been shown that BCL11B associates with NuRD complex (Cismasiu et al., 2005). Thus, BCL11B could participate in the assembly or the targeting the NRC complex to the repressed genes. Further biochemical and genomic analyses are required to elucidate the mechanisms specifying the assembly and gene targeting of NAC and NRC.

### Both NAC and NRC contribute to Notch biological functions

Based on biochemical and genomic data including ours, we propose that upon activation of Notch signaling pathway, ICN1 can assemble both NAC and NRC to directly activate and repress specific subset of genes. The integration of the two activities generates specific NOTCH outcomes allowing a tight regulation of Notch-mediated cell differentiation and proliferation. In support, we found that NRC target the B-cell lineage transcription factor PAX5 and the MYC repressor MAD4. PAX5 plays a central role in restricting the differentiation of lymphoid progenitors toward the B-cell lineage (Nutt et al., 1999; Urbanek et al., 1994). PAX5 (^-^/-) pro-B cells cultured upon NOTCH activation adopt a T and NK cell potential (Carotta et al., 2006; Hoflinger et al., 2004). Remarkably, bone marrow stromal cell lines that were engineered to express Notch ligands lose their capacity to support the differentiation of hematopoietic progenitors into B cell, and acquire the ability to produce mature T-cells (Carotta et al., 2006; Hoflinger et al., 2004; Jaleco et al., 2001; Schmitt and Zuniga-Pflucker, 2002). Notch-mediated suppression of B cell program was believed to results from an indirect mechanism (Jundt et al., 2008). Here we show that, in T-ALL cells, the NRC is bound to the CSL DNA binding motif within PAX5 locus to repress its transcription. Notch suppression in T-ALL results in loss of binding of the NRC subunits and transcriptional derepression of otherwise silent PAX5 and its target genes (CD79a and VPreB). Remarkably, deletion of the CSL binding motif within the PAX5 locus results in loss of Notch signaling mediated regulation of PAX5 transcription. Moreover, expression of ICN1 in the Pre-B cell leukemia cell line B-ALL (REH) not only repressed transcription of PAX5 and its target genes, but it also induced the expression of the T-cell associated transcription factors TCF-7. In light of our data, we propose that Notch-mediated T cell committement requires the induction of T cell program mediated by NAC and direct repression of B cell program by NRC.

Using gene expression profiling and chromatin immunoprecipitation approaches, MYC was identified as a direct and essential target of Notch signaling in T-ALL cell lines (Herranz and Ferrando, 2015; Margolin et al., 2009; Palomero et al., 2006; Weng et al., 2006). Importantly, MYC inhibition interfered with the proliferative effects of activated Notch, while retroviral expression of MYC could partially rescue GSI-induced growth arrest in some T-ALL cell lines (Weng et al., 2006). Consistent with these observations, several studies highlighted the importance of MYC in Notch-mediated normal and oncogenic functions (Chiang et al., 2016; Palomero et al., 2006; Ryan et al., 2017; Sharma et al., 2006; Wang et al., 2011; Weng et al., 2006; Yashiro-Ohtani et al., 2014). Our experiments show that upon Notch activation, the NRC is recruited to the promoter of MYC repressor MAD4 to block its expression. We find that inactivation of Notch pathway in T-ALL cells results in down modulation of MYC and up-regulation of MAD4 and MAX favoring the MAD-MAX heterodimer formation known to target and repress Myc-dependent transcription and proliferation. Of note, our attempt to further characterizes SUPT1 clones in which the CSL binding sites within the MAD4 locus was invalidated using CRISPR-CAS9 technology were unsuccessful due to the fact that the derived clones containing the mutation did not proliferate to generate sufficient amount of cells (data not shown). Interestingly, overexpression of MAD4 during hematopoietic development severely decreased early progenitor proliferation indeed (Boros et al., 2011). We propose that Notch-mediated regulation of Myc involves a cooperative action involving transcriptional activation of Myc by the NAC and transcriptional repression of MAD4 by the NRC. Thus, based on proteomic and genomic data, we propose that ICN1 can directly activate or repress target genes and that both activities contribute to its developmental and oncogenic functions. The integration of both Notch-activated and repressed genes should generate specific Notch outcomes, allowing an additional level of regulation downstream of ICN release that is central to the pleiotropic nature of this pathway.

## ACKNOWLEDGMENTS

We are thankful to Kourtis and Lazaris’s lab for providing us access to RNAseq before publication release. We thank Marion Helsmoortel for her precious help using CRISPR-CAS technology. Raoul Raffel for genomic data processing and Ségolène Augé for DNA motifs analyses. We are grateful to MGX platform for performing the ChIP-seq sequencing. This work was supported by “Institut National du Cancer INCA PLBIO14-344” to M.B. and J.D.L., “Fondation pour la Recherche Medicale FRM équipe labelisée” to M.B. and DEQ20160334940 to O.C. including support to A.H. and the Labex EpiGenMed « Investissements d’avenir » (ANR-10-LABX-12-01). D.D.was supported by MRT fellowship.

## AUTHOR CONTRIBUTIONS

M.B., C.M.D. Y.L., J.D.L. conceived the study. C.M.D. and A.Y. performed experiments. D.D. performed most of the bioinfo analyses. A.H. processed the Promoter Capture HI-C data. All the authors analyzed and discussed the data. M.B., C.M.D., O.C., wrote the manuscript with input from all authors.

The authors declare no competing interests.

## SUPPLEMENTAL INFORMATION

Supplemental information includes five figures.

## SUPPLEMENTAL FIGURE LEGENDS

**Figure S1.**
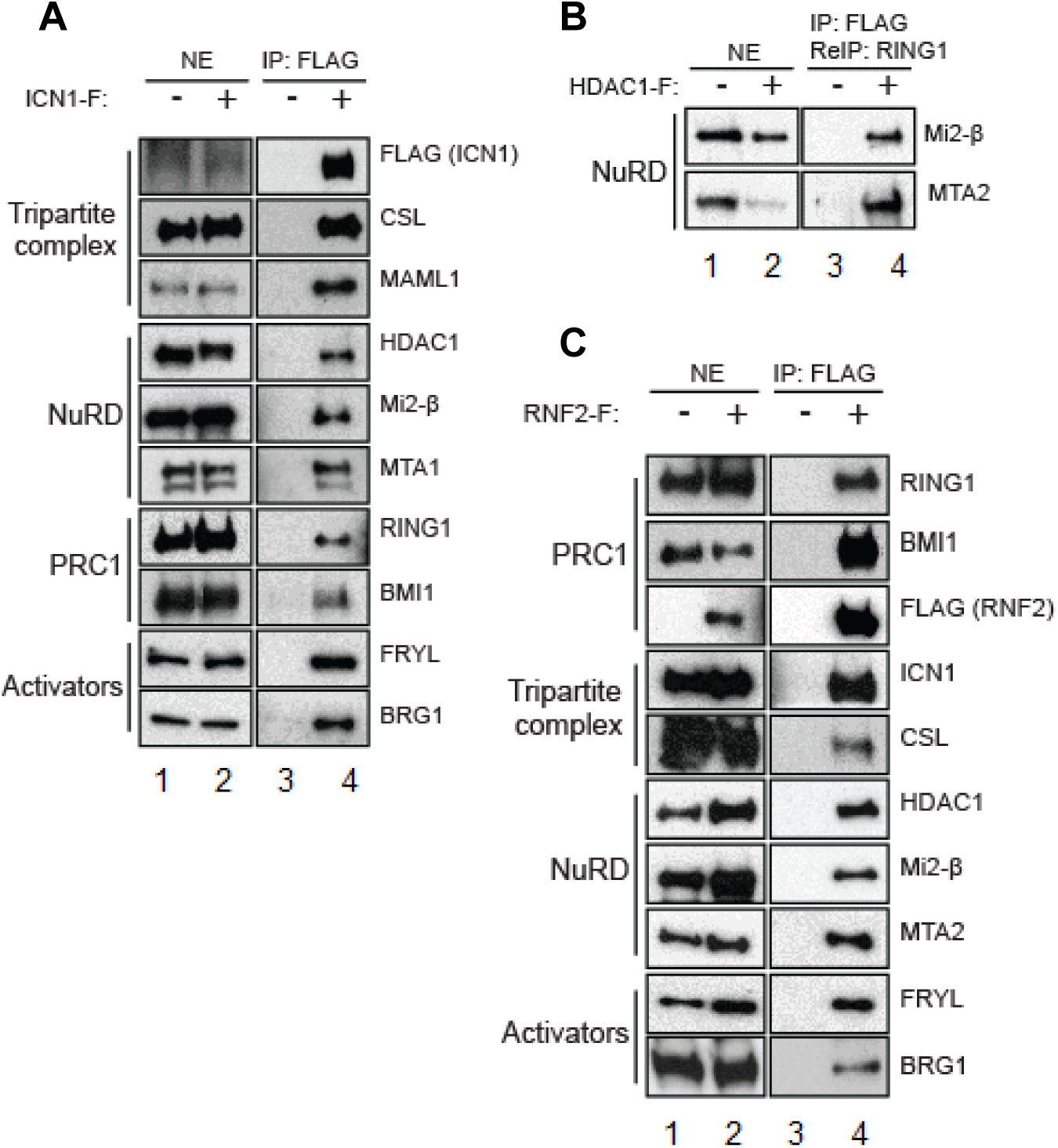
Biochemical characterization of ICN1 and its cofactors. Related to Figure 1. **(A)** Immunoprecipitation of ICN1-FLAG in SupT1 cells, stably expressing FLAG-tagged ICN1, analyzed by immunoblot. IP, immunoprecipitation; NE, nuclear extract. **(B)** Immunoprecipitation of HDAC1-FLAG followed by a second immunoprecipitation of RING1 (ReIP) in SupT1 cells, analyzed by immunoblot. **(C)** Immunoprecipitation of RNF2-FLAG in SupT1 cells, analyzed by immunoblot.

**Figure S2.**
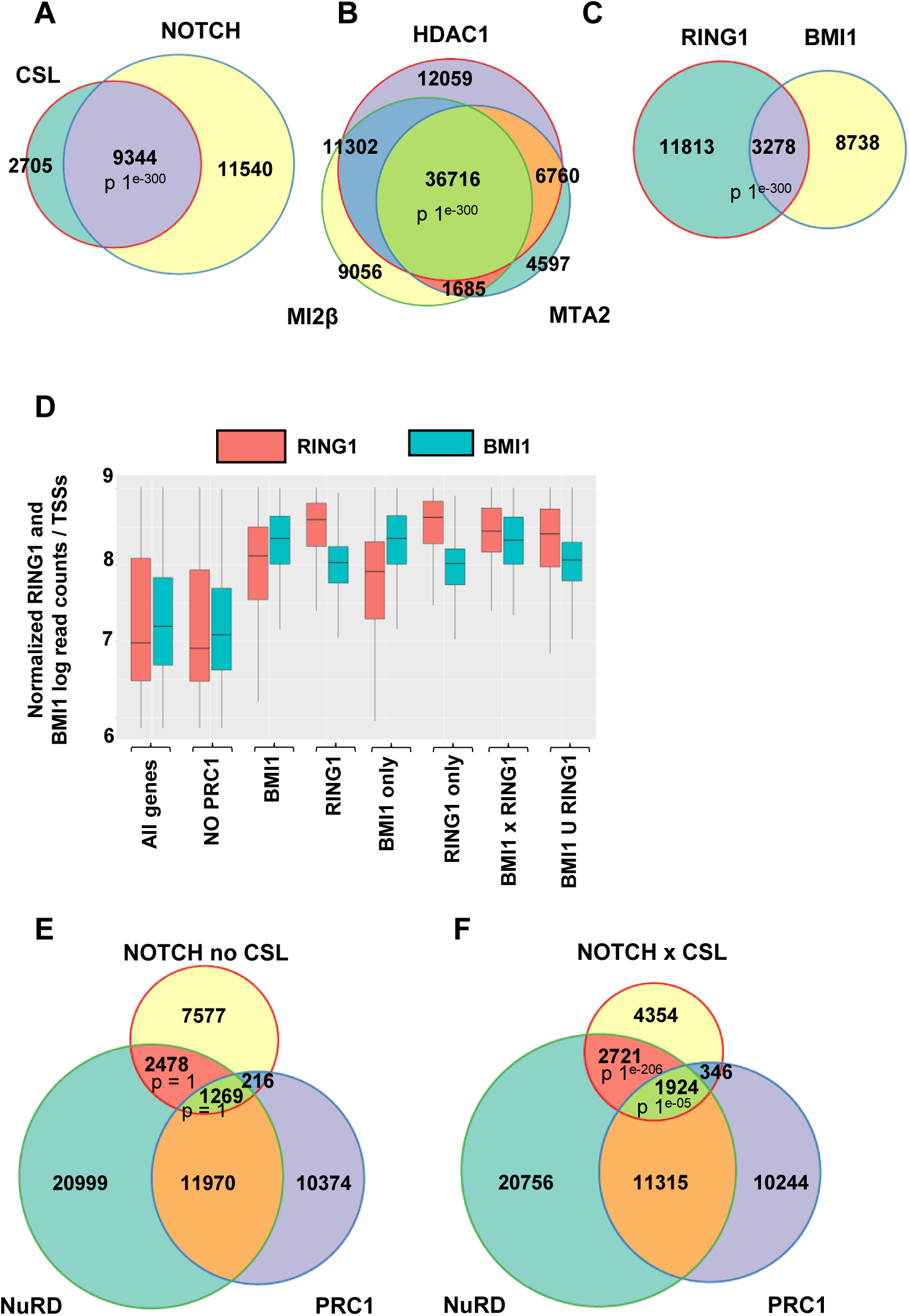
The NRC subunits overlap on chromatin in the presence of CSL., Related to Figure 2. **(A)** Venn diagrams showing the significant overlap among the genomic binding sites of Notch, and CSL. P-value was calculated using Fisher exact test. **(B)** Same as (A) with NuRD subunits: HDAC1, MI2β, and MTA2. **(C)** Same as (A) with PRC1 subunits: RING1 and BMI1. **(D)** Boxplot analysis showing the levels of BMI1 and RING1 binding on TSS (as normalized ChIP-seq counts). From left to right, boxplots represent RING1 (red) or BMI1 (blue) levels at all TSS (all genes) and TSS subgroups that harbor ‘no PRC1’, ‘BMI1’, ‘RING1’, ‘BMI1 only’, ‘RING1 only’, ‘intersection BMI1 and RING1’ and ‘union BMI1 and RING1’. Y-axis: Levels of normalized ChIP-seq reads. **(E)** Venn diagram showing the intersection between NOTCH, NuRD and PRC1 binding sites depending on NOTCH peaks excluding CSL. P-values were calculated using Fisher exact test. **(F)** Same as (F) on CSL-associated NOTCH peaks.

**Figure S3.**
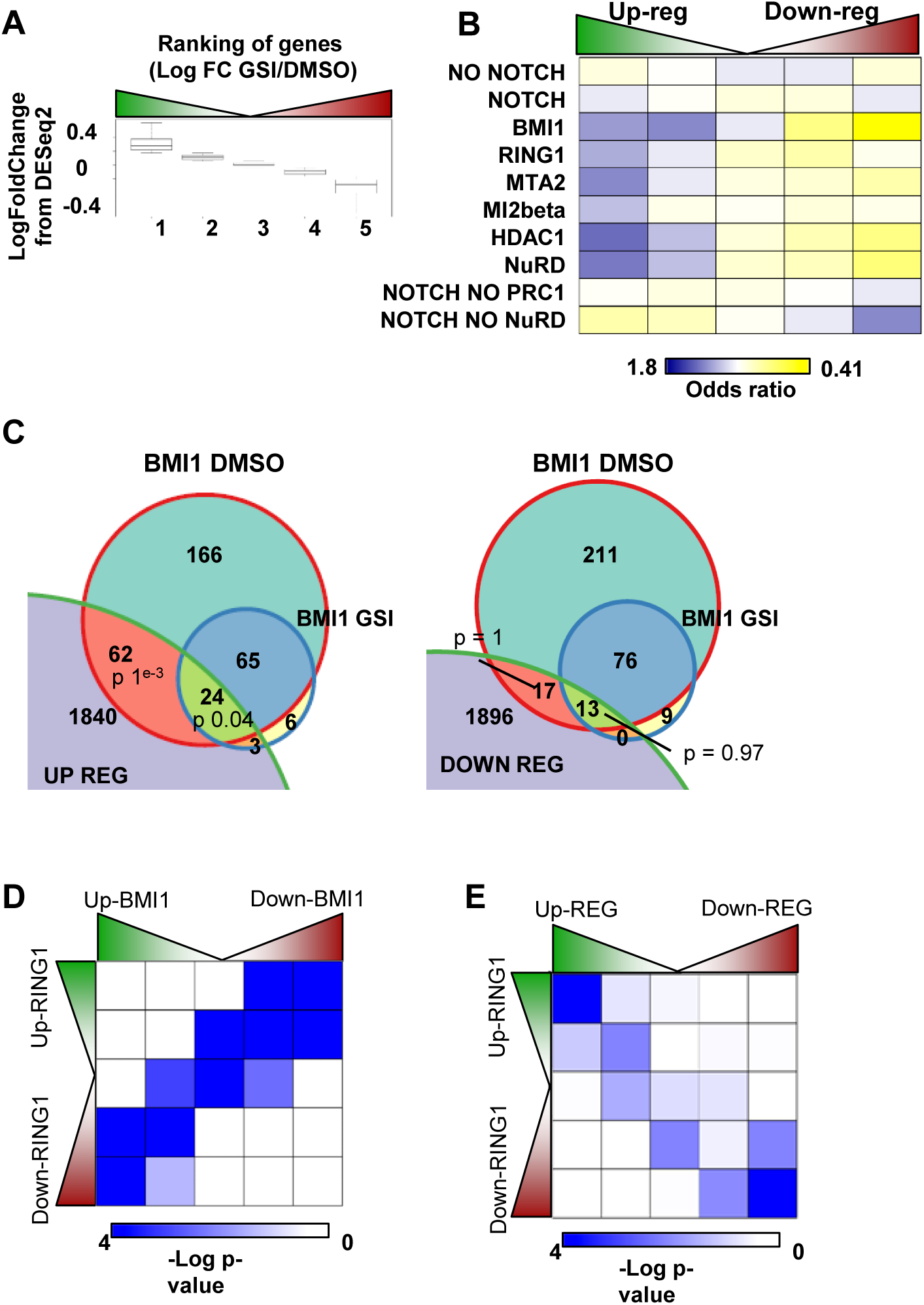
BMI1 and RING1 binding to NOTCH target genes is dynamically regulated, Related to Figure 3. **(A)** Boxplot showing the log fold change in gene expression as quantified by DEseq2 for RNAseq in GSI-treated as compared to DMSO-treated cells. **(B)** Same as Figure 3B, Shown in odds ratio. **(C)** Venn diagrams showing the intersection between the list of genes that are up-regulated or down-regulated upon GSI treatment depending on the persistence of BMI1 binding sites in DMSO compare to GSI conditions as detected by Macs2. P-values were calculated using Fisher exact test. **(D)** Intersection matrix showing that NOTCH depletion induces a dynamic RING1 binding that mirrors the BMI1 level. P-values were calculated using a Fisher exact test. **(E)** Intersection matrix showing that NOTCH depletion induces a dynamic RING1 binding depending on the expression level of NOTCH target genes. P-values were calculated using a Fisher exact test.

**Figure S4.**
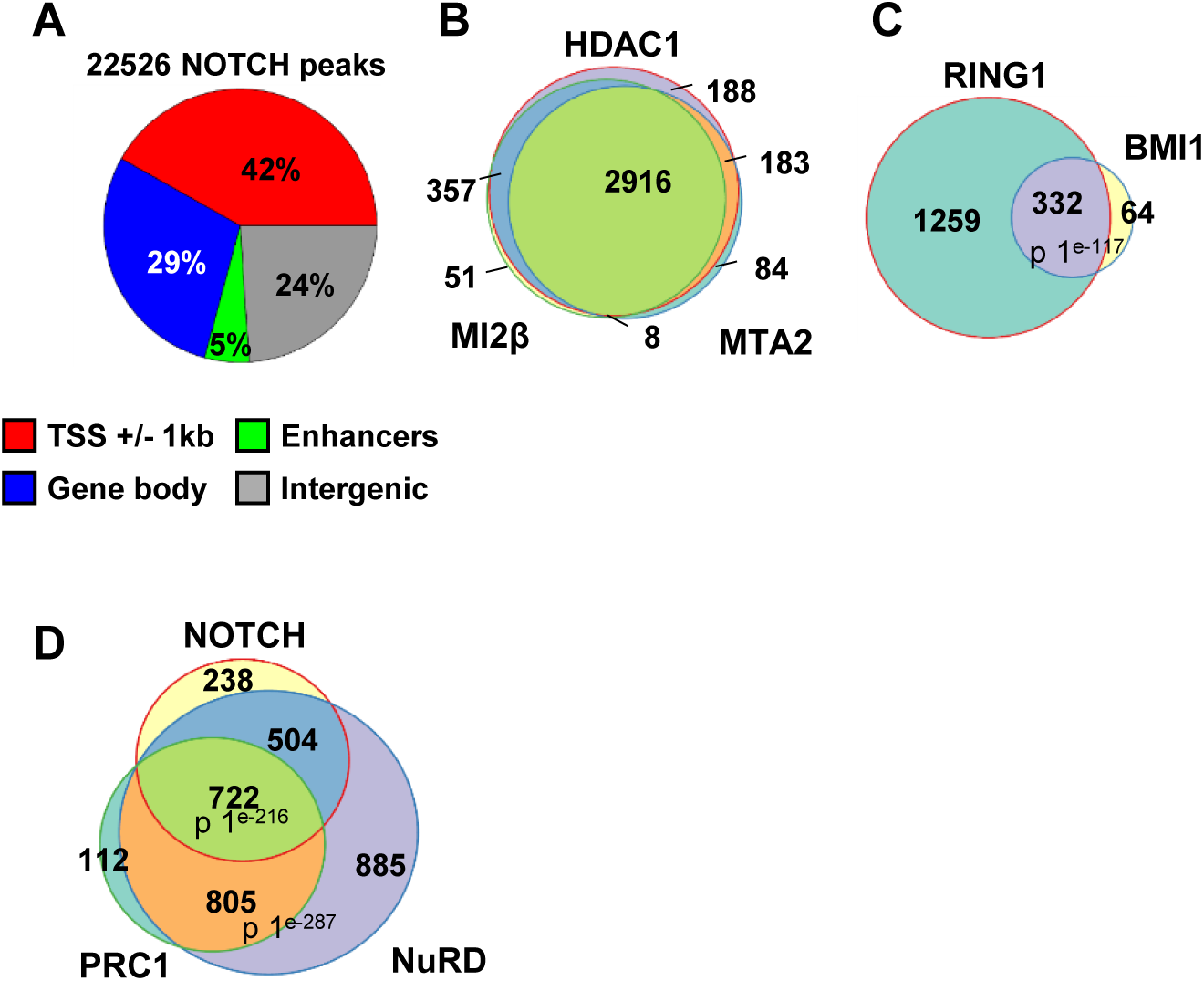
The NRC subunits co-localize on enhancers, Related to Figure 4. **(A)** Pie chart showing the relative distribution for NOTCH binding sites according to 4 sub-groups: TSSs (+/- 1 kbp), gene bodies (start to end), enhancers (as identified by the Fantom5 project) or in intergenic regions. **(B)** Venn diagram showing the co-binding of NuRD subunits, HDAC1, MTA2 and MI2β in the context of enhancers. P-values were calculated by Fisher exact test. **(C)** Same as (B) with PRC1 subunits, BMI1 and RING1. **(D)** Venn diagram showing the significant overlap among the genomic binding sites of Notch, NuRD, and PRC1 in the context of enhancers. ‘NuRD’ and ‘PRC1’ binding sites represent the overlapping sites where all NuRD (HDAC1, MI2β, and MTA2) and PRC1 (RING1 and BMI1) subunits were co-localized (See Fig.S4B-C). P-value was calculated using Fisher exact test.

**Figure S5.**
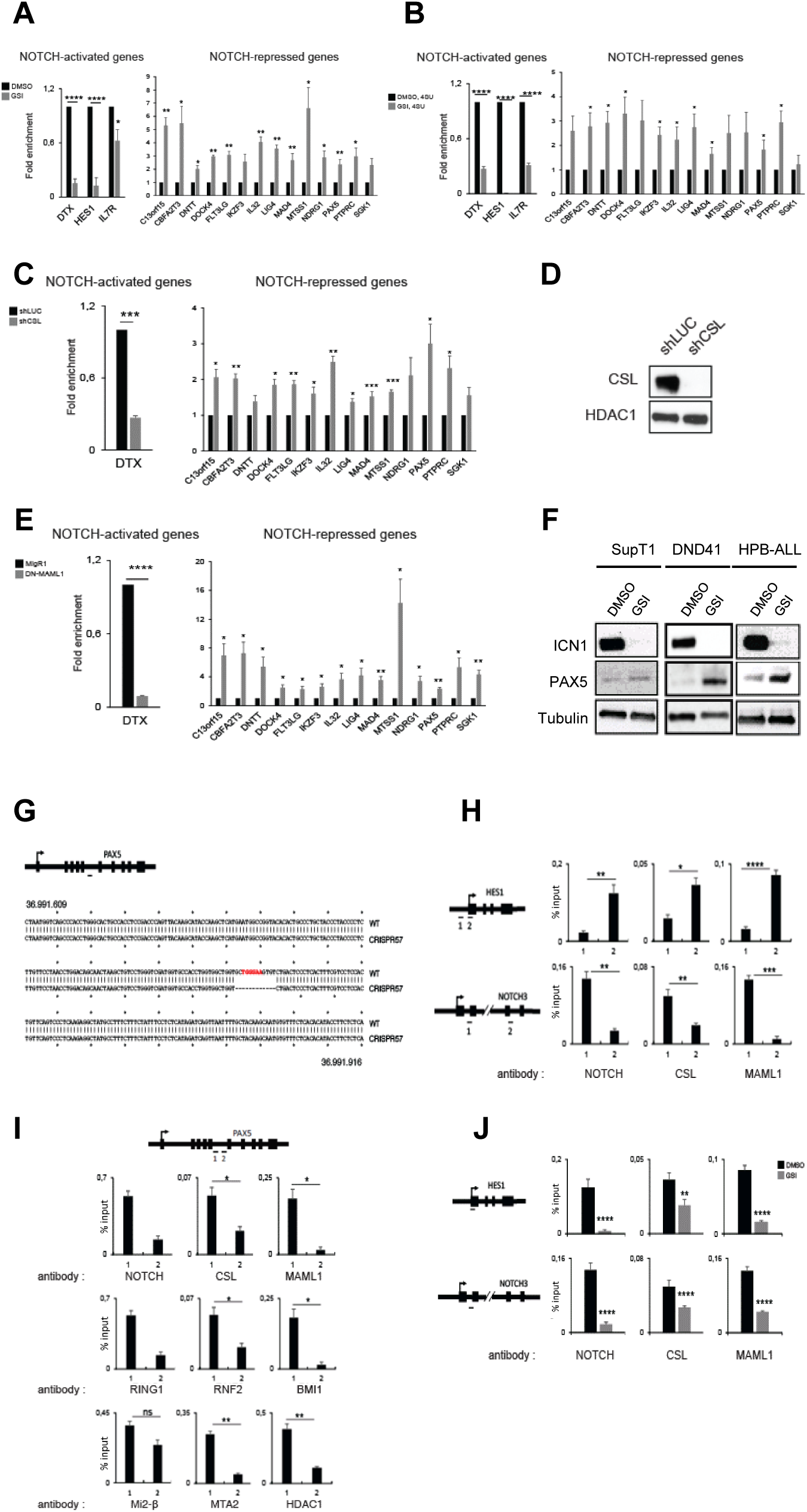
NOTCH signaling pathway transcriptionally represses NOTCH target genes through direct binding, Related to Figure 5. **(A)** qPCR measurement of NOTCH-target genes mRNA level inSupT1 cells treated 5 days with DMSO (black) or GSI (grey). **(B)** Analysis of the relative nascent mRNA expression of NOTCH-target genes by qRT-PCR in SupT1 cells treated 2 days with DMSO or GSI and incubated 15 min with 4sU component. **(C)** Relative mRNA expression of NOTCH-target genes in SupT1 cells treated 5 days with shLUC control or shCSL. mRNA levels were normalized to those of GAPDH. **(D)** The knockdown efficiency of CSL was monitored by immunoblotting of protein extracts from SupT1 cells. **(D)** Same as (C) using control or DN-MAML1. **(E)** Immunoblot analysis of protein extracts from SupT1, DND41, HPB-ALL cells. Cells were treated 5 days with DMSO or GSI. **(G)** Alignment of the *PAX5* genomic region (36.991.609-36.991.916) between wild-type and CRISPR57 SupT1 cell lines, the bold red letters correspond to CSL binding site deleted by CRISPR assay. The dark dash indicates the approximate position on PAX5 locus of the sgRNA used for the CRISPR experiment. **(H)** Locus occupancy analysis of NOTCH, CSL and MAML1 binding on HES1 and NOTCH3 by qChIP assay in SupT1 T-ALL cells. The amplicons are illustrated by black dashes. **(I)** Analysis by qChIP experiment of the binding of the NRC components on PAX5 locus in SupT1 T-ALL cells. The position of the amplicons is illustrated by black dashes. **(J)** Analysis of the locus occupancy of NOTCH, CSL and MAML1 on *HES1* and *NOTCH3* CSL-binding sites in SupT1 cells treated 2 days with DMSO or GSI. The position of the PCR amplicons is illustrated with black dashes.

## STAR METHODS

- **KEY RESOURCES TABLE** (**separately)**
- **CONTACT FOR REAGENT AND RESOURCE SHARING**
- **EXPERIMENTAL MODEL AND SUBJECT DETAILS**
  - Cells lines
  - Treatment of cell lines
- **METHOD DETAILS**
  - shRNA and expression vectors
  - Virus production and cell line transduction
  - Purification of complexes
  - Western Blot analysis
  - Quantitative RT-PCR and nascent transcripts
  - Chromatin Immunoprecipitation assays (ChIP) and library preparation
  - CRISPR-CAS9 experiment
- **QUANTIFICATION AND STATISTICAL ANALYSIS**
  - Statistical analyses if ChIP-Seq data
  - Statistical analyses of gene expression data
  - Enhancer-promoter interactions by integrating Promoter Capture Hi-C (PCHi-C) and the identification of T cell Enhancers
- **DATA AND SOFTWARE AVAILABILITY**

## KEY RESOURCES TABLE (separately)

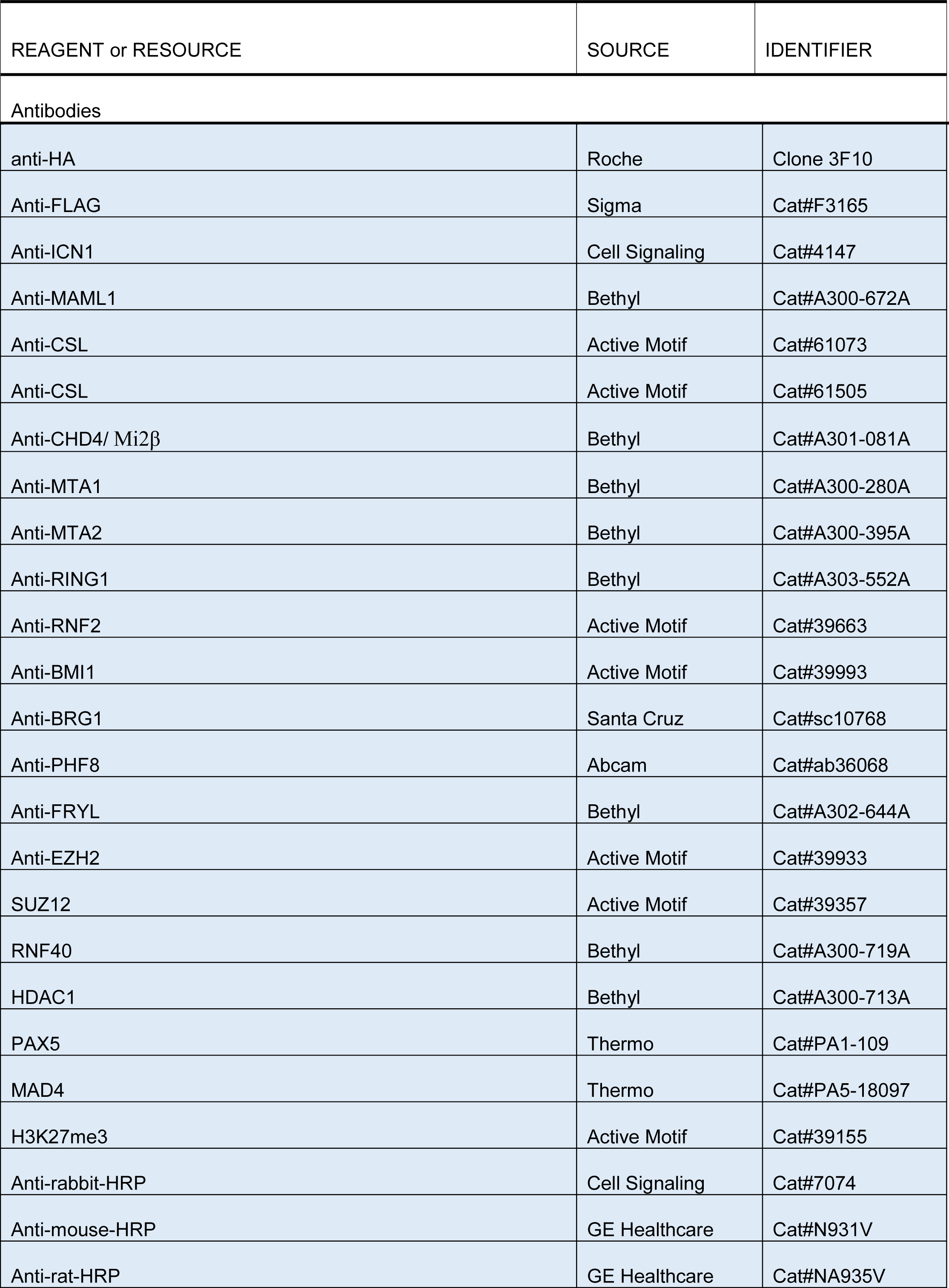
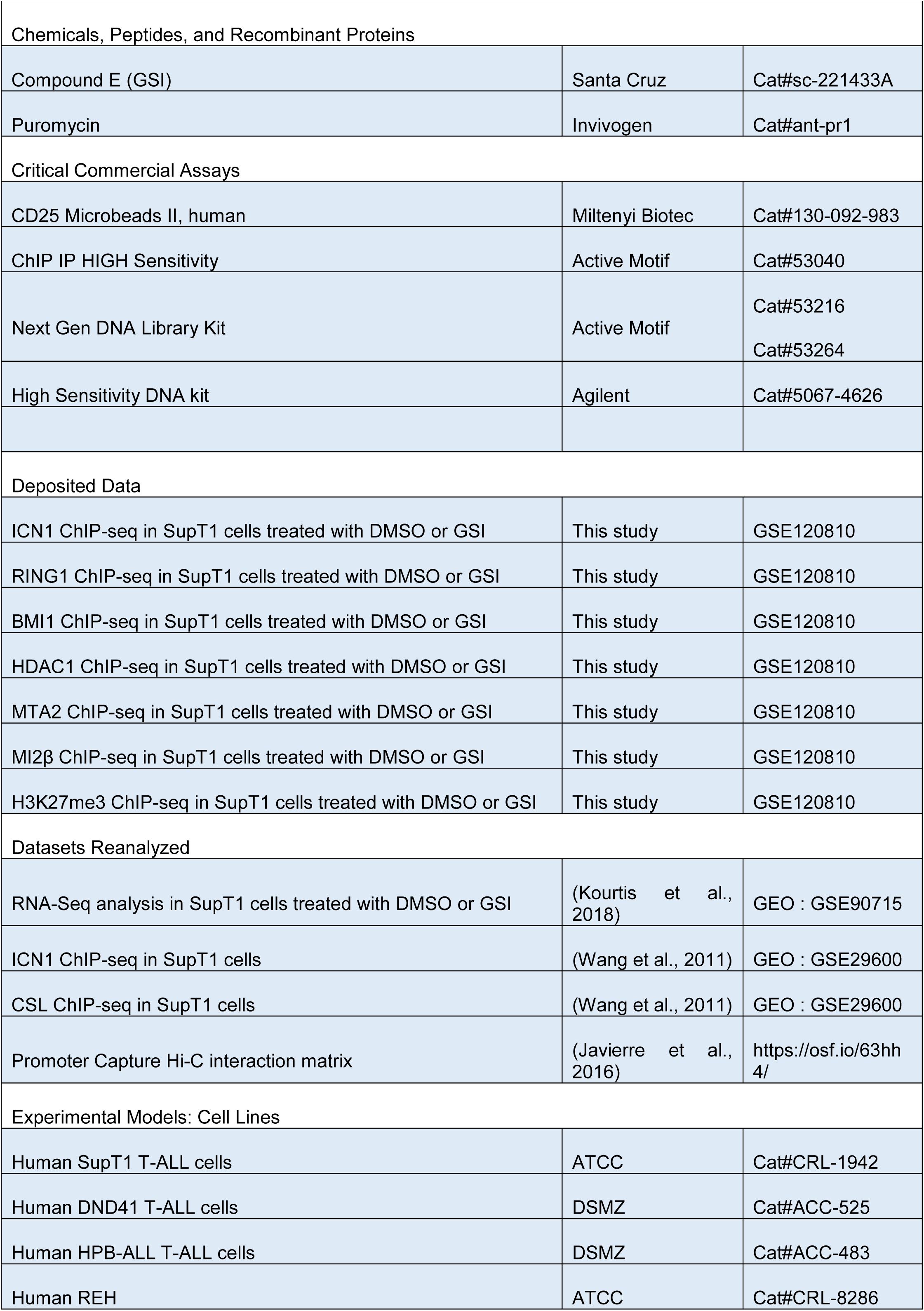
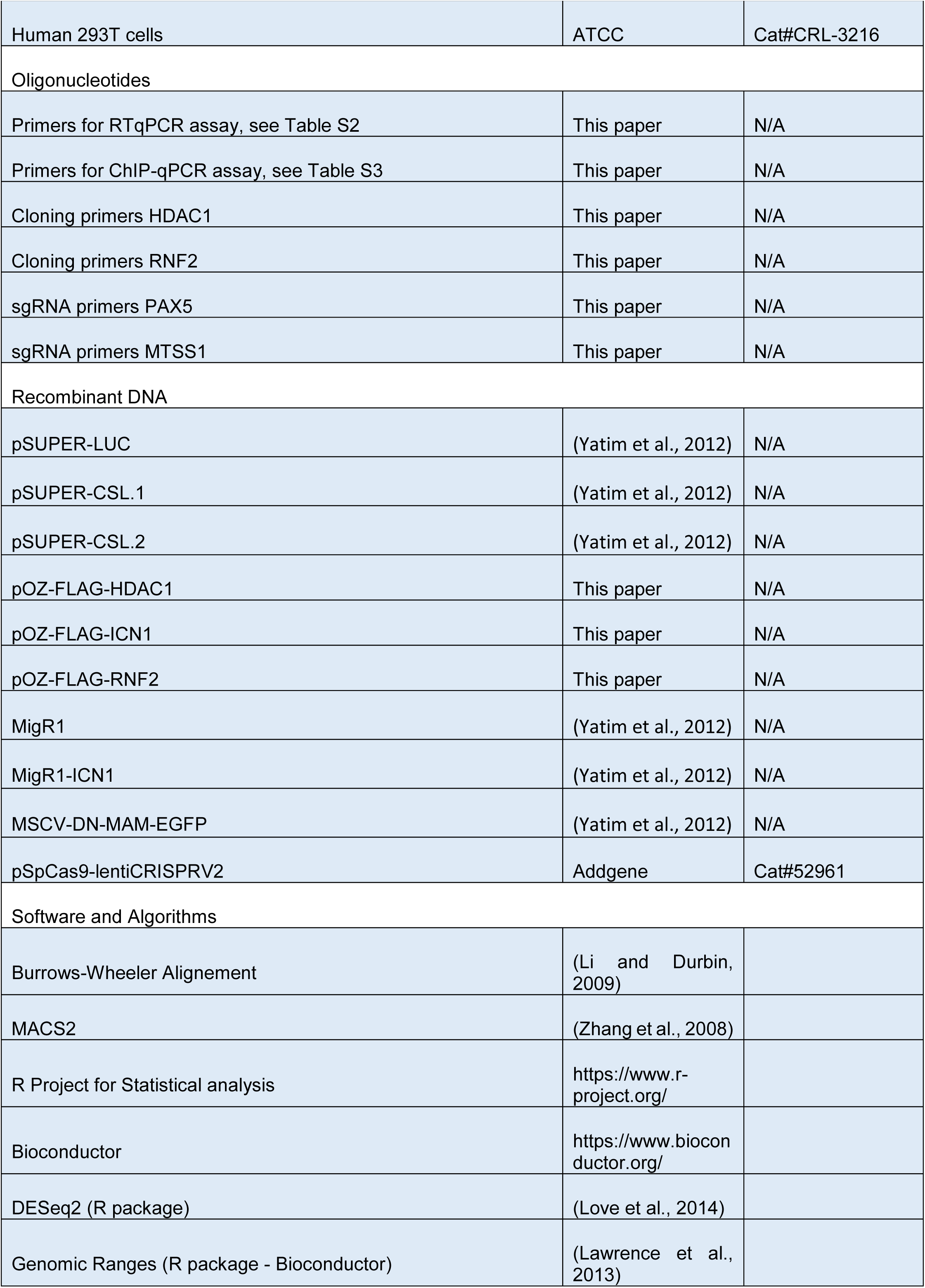
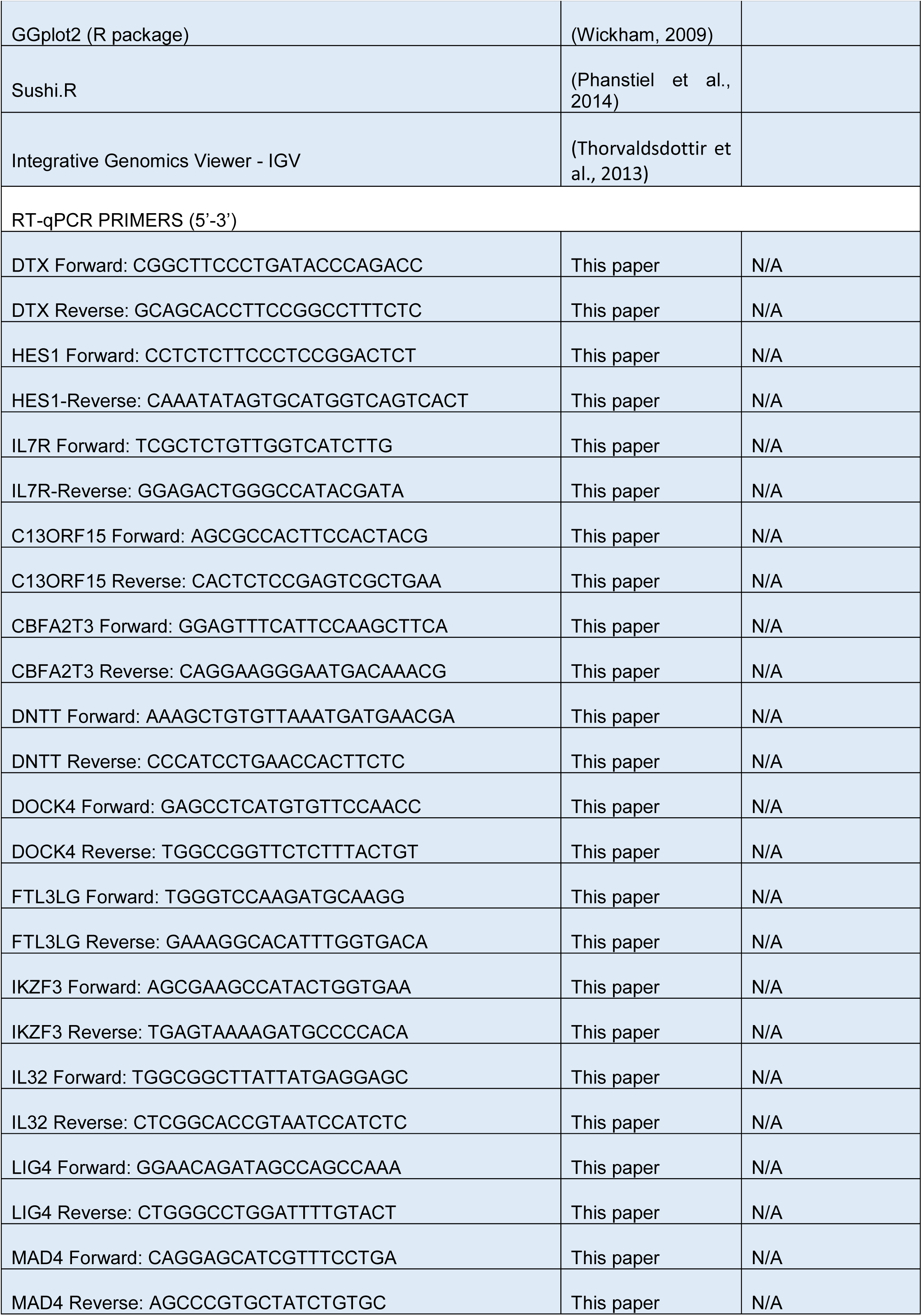
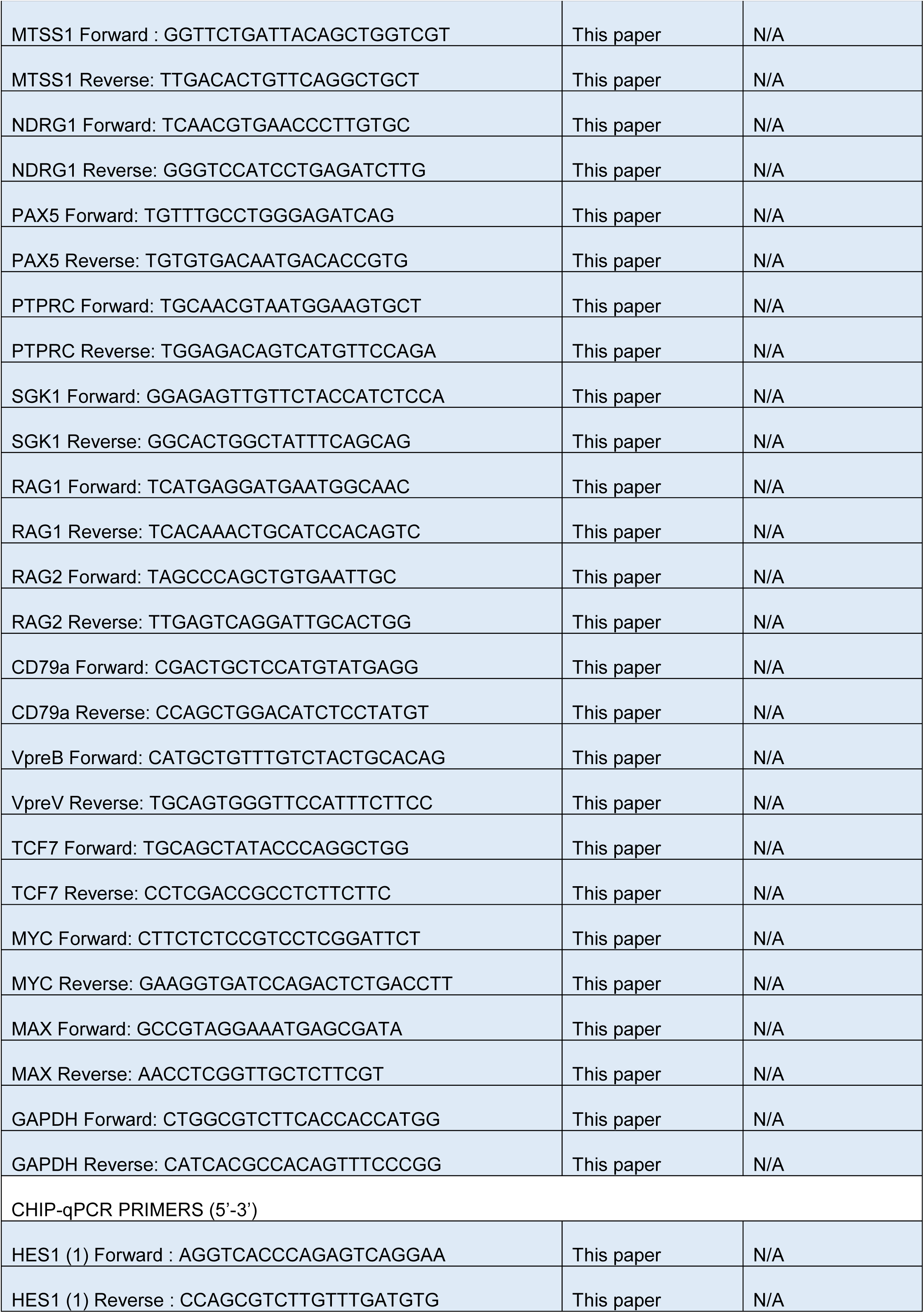
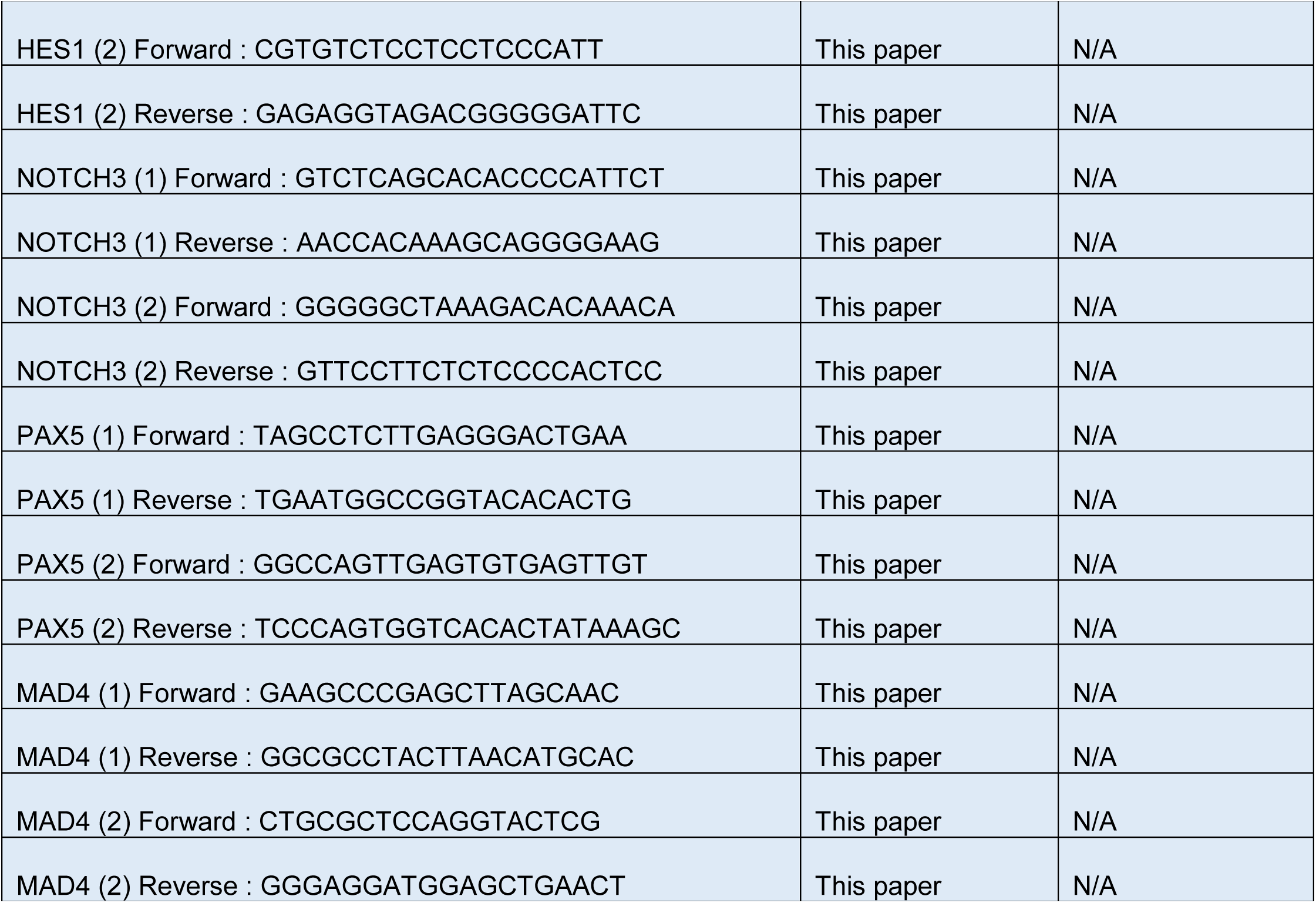

## CONTACT FOR REAGENT AND RESOURCE SHARING

Further information and requests for resources and reagents should be directed to and will be fulfilled by the Lead Contact, Monsef Benkirane.

## EXPERIMENTAL MODEL AND SUBJECT DETAILS

### Cells lines

The Human T-ALL cell lines SupT1, DND41, HPB-ALL were maintained in RPMI1640 Medium (Lonza) supplemented with 10% (v/v) heat-inactivated fetal calf serum (Lonza), 1000U.mL^-1^ penicillin/streptomycin (Lonza) and 20mM Ultraglutamine (Lonza). The Human 293T cells were maintained in DMEM Medium supplemented with 10% (v/v) heat-inactivated fetal calf serum (Lonza), 1000U.mL^-1^ penicillin/streptomycin (Lonza) and 20mM Ultraglutamine (Lonza).

### Treament of cell lines

Notch signaling was inhibited by treating T-ALL cells with the γ-secretase inhibitor (GSI) compound E at the final concentration of 1µM. For 2 days experiments, the T-ALL cells were treated on day 0 and 1. For 5 days experiment, T-ALL cells were treated on day 0, 1, 2 and 4.

## METHOD DETAILS

### shRNA and expression vectors

Overexpression in T-ALL cell lines were performed according to Yatim et al. using MigR1, MigR-ICN1 and MSCV-DN-MAM-EGFP, pOZ-ICN1-HA and pOZ-ICN1-FLAG. For HDAC1-FLAG and RNF2-FLAG expression, retroviral pOZ constructs containing a single tag (FLAG) were obtained by modifying the pOZ-Flag/HA (F/H) vector (Nakatani and Ogryzko, 2003) and the pOZ.puro-F/H vector (Kumar et al., 2009). Human HDAC1 and RNF2 were PCR amplified from SupT1 cDNA and inserted into the XhoI/NotI sites of pOZ vectors. All constructs were verified by sequencing. shRNA-mediated knockdown against CSL was performed according to Yatim et al. using pSUPER-LUC, pSUPER-CSL.1, pSUPER-CSL.2.

### Virus production and cell line transduction

293T cells were transfected with a packaging mixture and the retroviral vector (pOZ, pSUPER) using the calcium phosphate precipitation method. For transfection, 5µg of the retroviral vector, 2.5µg of the packaging plasmid (gag/pol) and 2.5µg of the envelope plasmid were mixed with 100µL of CaCl2 (1.25M) and 500µL of HBS2X in a final volume of 1mL. The mixture was incubated 1min at room temperature then added dropwise to the cells. The medium was changed the following day and the viral-containing supernatant was collected 24 hours after transfection, filtered through a 0.45 µm filter and subsequently used to infect cells. To establish stable SupT1 cell lines expressing tagged FLAG-HDAC1 and HA-ICN1 or FLAG-RNF2, we first transduced SupT1 with recombinant retroviruses encoding for FLAG-HDAC1 and puromycin resistance marker. Transduced cells were selected by puromycin treatment (2µg/mL). Second, we transduced SupT1 expressing FLAG-HDAC1 with recombinant retroviruses encoding for HA-ICN1 and IL-2 receptor subunit alpha resistance gene. Transduced cells were purified by affinity using CD25 MicroBeads II (Miltenyi Biotec 130-092-983). For shRNA-mediated knockdown experiments, cells were transduced with recombinant retroviruses or lentiviruses. After an overnight incubation, a second round and a third round of infection was performed using the same vector (or a second shRNA targeting the same mRNA (for CSL)) (Yatim et al., 2012). The medium was refreshed the following day and puromycin was added 72 hours post-infection at a final concentration of 2µg/mL. Protein expression was analyzed by western blot after 5 days of selection. All the experiments were performed at 5 days post-transduction.

### Purification of complexes

Nuclear extracts were prepared using the Dignam protocol with slight modifications (Dignam et al., 1983). For the purification of HDAC1-ICN1-associated complexes, SupT1 cells stably expressing Flag tagged HDAC1 and HA tagged ICN1 and control SupT1 were harvested by centrifugation, washed in cold PBS and resuspended in 4 packed cell pellet volumes of hypotonic buffer (20 mM Tris-HCl pH 7.4, 10 mM NaCl, and 1.5 mM MgCl2). The suspension was incubated on ice for 10 min and then cells were lysed by 12 strokes using a Dounce homogenizer fitted with a B pestle. The nuclei were pelleted by centrifugation and resuspended in one packed nuclear pellet volume of a buffer containing 20 mM Tris-HCl pH 7.4, 300 mM NaCl, 25% glycerol, 0.2 mM EDTA, 1.5 mM MgCl2 and PMSF. One packed nuclear pellet volume of a high salt buffer (containing 20 mM Tris-HCl pH 7.4, 720 mM NaCl, 25% glycerol, 0.2 mM EDTA, 1.5 mM MgCl2 and PMSF) was added dropwise to the suspension gently stirring with a magnetic bar. After stirring for 30 min at 4°C, the suspension was centrifuged at 13.000g for 30 min at 4°C and the supernatant was dialyzed against 100 volumes of buffer BC100 (20 mM Tris-HCl pH 7.4, 100 mM NaCl, 10% glycerol, 0.2 mM EDTA, 1.5 mM MgCl2 and PMSF) for 4 hours. The dialysate (nuclear extract) was cleared by centrifugation at 13.000g for 30 min. Nuclear extracts were incubated for 4 hr (at 4°C with rotation) with anti-FLAG M2 agarose beads (Sigma) (1% v/v) equilibrated in BC100. Beads were washed 3 times with 10 mL buffer B015 (20 mM Tris-HCl pH 7.4, 150 mM NaCl, 10% glycerol, 0.5 mM EDTA, 5 mM MgCl2, 0.05% Triton X-100, 0.1% Tween, and PMSF) and bound proteins were eluted with 4 bead volumes of B015 containing 0.2 mg/mL of FLAG peptide (Sigma) for 1 hr. The FLAG affinity purified complexes were further immunopurified by affinity chromatography using 10 µL of anti-HA conjugated agarose beads (Santa Cruz). After incubation for 4 hr, HA beads were washed 4 times with 800µL of buffer B015 in spin columns (Pierce, 69702) and eluted under native conditions using HA peptide (Roche). To isolate HDAC1-RING1 associated proteins, two-step affinity purification was performed on nuclear extracts from SupT1 cells stably expressing FLAG tagged HDAC1: a first IP-FLAG followed by a ReIP using RING1 antibody and protein G Sepharose beads (Fast flow, Sigma).

HDAC1-, ICN1- or RNF2-associated proteins were isolated through one-step affinity purification on nuclear extracts from SupT1 cells stably expressing FLAG tagged HDAC1/ICN1/RNF2. For all the immunoprecipitation experiments, the eluted proteins were resolved by western blot analysis using indicated antibodies.

### Western Blot analysis

Cells were lysed in lysis buffer (50 mM Tris-HCl, 120 mM NaCl, 5 mM EDTA, 0.5% NP-40 and PMSF) and briefly sonicated. Cell lysates and immunoprecipitates were boiled in LDS sample buffer and resolved on a 4-15% SDS-PAGE gel (Biorad), in transfer buffer (20% methanol, 25 mM Tris, 192 mM Glycine, 0.037% SDS) during 90 min at 100V. Proteins were liquid-transferred to nitrocellulose membrane. Primary antibodies were incubated overnight at room temperature in PBS 1% milk 0.1% tween; secondary antibodies were incubated 1h at room temperature in PBS 5% milk 0.1% tween.. Secondary antibodies are used at 1: 2000: anti-rabbit-HRP, anti-mouse-HRP and anti-rat-HRP.

### Quantitative RT-PCR and nascent transcripts

RNAs were isolated using the Trizol reagent (Invitrogen) and reverse transcription was performed with 200ng to 1µg of RNA using SuperScript IV (Invitrogen) and oligodT per the manufacturer’s instructions. PCR measurements were performed in triplicate using Master Mix SYBR Green (Roche). Amplification was carried out in the LightCycler480 (Roche). The average of the technical replicates was normalized to GAPDH levels using the comparative CT method (2-ΔΔCT). Averages and standard deviations of at least 3 experiments are shown in the figures. RT-qPCRs were performed using the indicated primers (Key Resources Table). For the analysis of nascent mRNA, total RNA was isolated using Trizol reagent (Invitrogen). 4sU labeled mRNA was purified using biotin-streptavidin assay (Schwalb et al., 2016) and reverse transcription was performed as described above.

### Chromatin Immunoprecipitation assays (ChIP) and library preparation

ChIP experiments were performed in SupT1 T-ALL cells using the ChIP IP High Sensitivity (Active Motif 53040). ChIP DNA was analyzed by qPCR using Master Mix SYBR Green (Roche) using specific primers as described in Key Resources Table. Amplification was carried out in the LightCycler480 (Roche). PCR measurements were performed in duplicate. The average of technical replicates was normalized to GAPDH level using the comparative CT method. Averages and standard deviations of three experiments are shown in the figures. Antibodies used for ChIP experiment are indicated in Table S1.

ChIP-seq librairies were constructed using the Next Gen DNA Library Kit (Active Motif 53216 and 53264). Library quality was assessed using Agilent 2100 Bioanalyzer and Agilent High Sensitivity DNA assay. Sequencing was realized using HiSeq 2500 Illumina by Sequence By Synthesis technique. Image analysis and base calling were performed using the HiSeq Control Software and Real-Time Analysis component. De-multiplexing was performed using blc2fastq (Illumina). Data quality was assessed using Illumina software Sequencing Analysis Viewer and FastQC from the Babraham Institute.

### CRISPR-CAS9 experiment

sgRNA were designed, hybridized and cloned into pSpCas9-lentiCRISPRV2 (lentiCRISPR v2 was a gift from Feng Zhang (Addgene plasmid # 52961)) (according to (Ran et al., 2013; Sanjana et al., 2014)). Sequences of sgRNAs: sgPAX5: GCCACCTGGTGGCTGGTGCT; sg1. Production of lentiviral particules was realized as described above. SupT1 cells were transduced with lentiCRISPRV2-sgPAX5 particules. Two days post-puromycin selection, the cells were single cell selected into 96 wells plate. gDNA from several clones was PCR amplified by covering the region of interest and subsequently cloned (Invitrogen, Zero Blunt TOPO PCR cloning). The PCR clones were next sequenced and the CRISPR57 clone was selected as positive clone

## QUANTIFICATION AND STATISTICAL ANALYSIS

### Statistical analyses of ChIP-Seq data

After quality control using fastQC tool, ChIP-seq reads of ICN1, CSL, BMI1, RING1, HDAC1, MTA2, MI2β, and H3K27me3 in both condition DMSO and GSI have been aligned on hg38 reference genome using Burrows-Wheeler Aligner (BWA, http://bio-bwa.sourceforge.net/) and then normalized on the corresponding input. ChIP-Seq peaks of ICN1, CSL, BMI1, RING1, HDAC1, MTA2,MI2β, and H3K27me3 in both condition DMSO and GSI were identified using MACS2 with normalisation to the corresponding input sequenced in parallel. For peak calling analysis, ChIP-seq of ICN1 and CSL are obtained from geodataset GSE29600, all others ChIP-seq data, including ICN1 in DMSO and GSI conditions were generated in M. Benkirane team. All following analysis were done with R 3.4.2 version. Genes associated with peaks were identified by overlapping within the +/-500bp region around TSS. Enhancers associated with peaks were identified by overlapping peaks summit with each enhancer range.

Overlapping analyses were performed using « GenomicRanges » R functions (https://bioconductor.org/packages/release/bioc/html/GenomicRanges.html) as previously done (Stadelmayer et al., 2014). Enrichments in Venn diagram and enrichment matrix were performed using a Fisher exact test, p-values are one-sided to reflect only enrichment (i.e. positive Odds ratio, alternative = « greater » of R Fisher. Test). Proportional Venn diagram were plotted with « Vennerable » R package (https://github.com/js229/Vennerable). Reads quantification were done for TSS over the +/-500bp region; for enhancers over the entire region of the enhancers and then normalized over enhancers length. Changes estimation in numbers of reads in boxplot was performed using a pair-wise Wilcoxon test comparing variation upon same subsets of genes. Scatter plot and boxplots have been done using ggplot2 R package functions (https://cran.r-project.org/web/packages/ggplot2/index.html). Binding dynamics of ICN1, RING1, BMI1, were performed in cells treated with GSI and DMSO for 24hrs. Binding was then analyzed by ranking genes according to the indicated (Notch or RING1) normalized ChIP-seq read counts in TSS +/-1kbp and ChIP-seq reads of ICN1, RING1, BMI1 were aligned by TSSs (position 0 with genes oriented towards the right). Analyses of enhancers-promoters association had been determined using promoter capture experiment and plots were obtained using adapted functions of Sushi R/Bioconductor package (https://bioconductor.org/packages/release/bioc/html/Sushi.html). Motif conservation analysis add been performed using functions from « Biostrings » R package.

### Statistical analyses of gene expression data

Gene expression analyses were first performed with microarray data. For genome-wide detection, we then integrated polyA+ RNAseq analysis from public dataset GSE90715 (Kourtis et al., 2018) using T-ALL cell line treated with DMSO or GSI for the same time (24 hrs). Gene expression levels in duplicates normalized by fpkm were directly recovered in DMSO and GSI conditions. After filtering low reads genes using R package HTSfilter, a differential analysis had been performed with DESeq2 R package for comparing expression level of each gene in GSI vs DMSO conditions. Up-regulated and down-regulated genes subsets were defined according to a p-value threshold <0,05 and a LogFoldChange threshold > 1 or < 1 respectively. Enrichments test between differential expressed genes and others subsets of genes such as genes associated with peaks were done using Fisher Exact test.

### Enhancer-promoter interactions by integrating Promoter Capture Hi-C (PPCHi-C) and the identification of T-cell Enhancers

PC Hi-C interaction matrix was downloaded from https://osf.io/63hh4/ (Javierre et al., 2016). From this matrix, both activated and non-activated CD4^+^ T cells interactions could be filtered out considering only those who had a high-confidence PCHi-C interactions (CHiCAGO score > = 5) as previously done by the authors. The identification of enhancers was originally performed by Ienasescu and colleagues (http://slidebase.binf.ku.dk) for differentially expressed T cells enhancers (Ienasescu et al., 2016) as part of the Fantom5 project (Lizio et al., 2015). We used high-confidence PCHi-C interactions (see PCHi-C method; (Javierre et al., 2016)), to generate an enhancer-promoter interaction matrix. Promoters (from the same genome release TxDb.Hsapiens.UCSC.hg19.knownGene R library) and T cells enhancers were first associated with left and right PCHi-C interacting fragments as defined by HindIII cutting sites. Then an association was performed to generate couples of enhancers and promoters for an estimation of their interaction between the corresponding fragments.

Further illustration of interaction (Figure 4) was performed using Sushiplot R/Bioconductor package (Phanstiel et al., 2014). The plotBedpe function was used to plot PCHiC interactions and two modifications were applied to plotGenes and plotBed. plotGenes function was modified for illustration legibility by avoiding gene overcrowding when plotting genomic region of interest (closeby genes < 1000 bp from the gene of interest were deleted). For the same readability reasons, modifications were brought to PlotBed function to allow clear representation of HindIII fragments position

## DATA AND SOFTWARE AVAILABILITY

### Data Resources

The accession number for the super series of data deposited to GEO and pertaining to this paper is GSE120810.

## REFERENCES

Aifantis, I., Raetz, E., and Buonamici, S. (2008). Molecular pathogenesis of T-cell leukaemia and lymphoma. Nat Rev Immunol 8, 380–390.

Andersson, E.R., Sandberg, R., and Lendahl, U. (2011). Notch signaling: simplicity in design, versatility in function. Development 138, 3593–3612.

Antila, C.J.M., Rraklli, V., Blomster, H.A., Dahlstrom, K.M., Salminen, T.A., Holmberg, J., Sistonen, L., and Sahlgren, C. (2018). Sumoylation of Notch1 represses its target gene expression during cell stress. Cell Death Differ 25, 600–615.

Ayer, D.E., and Eisenman, R.N. (1993). A switch from Myc:Max to Mad:Max heterocomplexes accompanies monocyte/macrophage differentiation. Genes Dev 7, 2110–2119.

Borggrefe, T., and Liefke, R. (2012). Fine-tuning of the intracellular canonical Notch signaling pathway. Cell Cycle 11, 264–276.

Bornelov, S., Reynolds, N., Xenophontos, M., Gharbi, S., Johnstone, E., Floyd, R., Ralser, M., Signolet, J., Loos, R., Dietmann, S., et al. (2018). The Nucleosome Remodeling and Deacetylation Complex Modulates Chromatin Structure at Sites of Active Transcription to Fine-Tune Gene Expression. Mol Cell 71, 56–72 e54.

Boros, K., Lacaud, G., and Kouskoff, V. (2011). The transcription factor Mxd4 controls the proliferation of the first blood precursors at the onset of hematopoietic development in vitro. Exp Hematol 39, 1090–1100.

Bray, S., Musisi, H., and Bienz, M. (2005). Bre1 is required for Notch signaling and histone modification. Dev Cell 8, 279–286.

Bray, S.J. (2006). Notch signalling: a simple pathway becomes complex. Nat Rev Mol Cell Biol 7, 678–689.

Busslinger, M. (2004). Transcriptional control of early B cell development. Annu Rev Immunol 22, 55–79.

Carotta, S., Brady, J., Wu, L., and Nutt, S.L. (2006). Transient Notch signaling induces NK cell potential in Pax5-deficient pro-B cells. Eur J Immunol 36, 3294–3304.

Chiang, M.Y., Wang, Q., Gormley, A.C., Stein, S.J., Xu, L., Shestova, O., Aster, J.C., and Pear, W.S. (2016). High selective pressure for Notch1 mutations that induce Myc in T-cell acute lymphoblastic leukemia. Blood 128, 2229–2240.

Cismasiu, V.B., Adamo, K., Gecewicz, J., Duque, J., Lin, Q., and Avram, D. (2005). BCL11B functionally associates with the NuRD complex in T lymphocytes to repress targeted promoter. Oncogene 24, 6753–6764.

Collins, K.J., Yuan, Z., and Kovall, R.A. (2014). Structure and function of the CSL-KyoT2 corepressor complex: a negative regulator of Notch signaling. Structure 22, 70–81.

de Pooter, R.F., Schmitt, T.M., de la Pompa, J.L., Fujiwara, Y., Orkin, S.H., and Zuniga-Pflucker, J.C. (2006). Notch signaling requires GATA-2 to inhibit myelopoiesis from embryonic stem cells and primary hemopoietic progenitors. J Immunol 176, 5267–5275.

Dege, C., and Hagman, J. (2014). Mi-2/NuRD chromatin remodeling complexes regulate B and T-lymphocyte development and function. Immunol Rev 261, 126–140.

Fortini, M.E. (2009). Notch signaling: the core pathway and its posttranslational regulation. Dev Cell 16, 633–647.

Freund, P., Kerenyi, M.A., Hager, M., Wagner, T., Wingelhofer, B., Pham, H.T.T., Elabd, M., Han, X., Valent, P., Gouilleux, F., et al. (2017). O-GlcNAcylation of STAT5 controls tyrosine phosphorylation and oncogenic transcription in STAT5-dependent malignancies. Leukemia 31, 2132–2142.

Fryer, C.J., White, J.B., and Jones, K.A. (2004). Mastermind recruits CycC:CDK8 to phosphorylate the Notch ICD and coordinate activation with turnover. Mol Cell 16, 509–520.

Gao, Z., Zhang, J., Bonasio, R., Strino, F., Sawai, A., Parisi, F., Kluger, Y., and Reinberg, D. (2012). PCGF homologs, CBX proteins, and RYBP define functionally distinct PRC1 family complexes. Mol Cell 45, 344–356.

Grandori, C., Cowley, S.M., James, L.P., and Eisenman, R.N. (2000). The Myc/Max/Mad network and the transcriptional control of cell behavior. Annu Rev Cell Dev Biol 16, 653–699.

Grinberg, A.V., Hu, C.D., and Kerppola, T.K. (2004). Visualization of Myc/Max/Mad family dimers and the competition for dimerization in living cells. Mol Cell Biol 24, 4294–4308.

Guruharsha, K.G., Kankel, M.W., and Artavanis-Tsakonas, S. (2012). The Notch signalling system: recent insights into the complexity of a conserved pathway. Nat Rev Genet 13, 654–666.

Han, X., Ranganathan, P., Tzimas, C., Weaver, K.L., Jin, K., Astudillo, L., Zhou, W., Zhu, X., Li, B., Robbins, D.J., et al. (2017). Notch Represses Transcription by PRC2 Recruitment to the Ternary Complex. Mol Cancer Res 15, 1173–1183.

Herranz, D., and Ferrando, A.A. (2015). An oncogenic enhancer enemy (N-Me) in T-ALL. Cell Cycle 14, 167–168.

Hoflinger, S., Kesavan, K., Fuxa, M., Hutter, C., Heavey, B., Radtke, F., and Busslinger, M. (2004). Analysis of Notch1 function by in vitro T cell differentiation of Pax5 mutant lymphoid progenitors. J Immunol 173, 3935–3944.

Huang, X., Huang, B., and Li, H. (2009). A fast level set method for synthetic aperture radar ocean image segmentation. Sensors (Basel) 9, 814–829.

Hutchins, A.S., Mullen, A.C., Lee, H.W., Sykes, K.J., High, F.A., Hendrich, B.D., Bird, A.P., and Reiner, S.L. (2002). Gene silencing quantitatively controls the function of a developmental trans-activator. Mol Cell 10, 81–91.

Jaleco, A.C., Neves, H., Hooijberg, E., Gameiro, P., Clode, N., Haury, M., Henrique, D., and Parreira, L. (2001). Differential effects of Notch ligands Delta-1 and Jagged-1 in human lymphoid differentiation. J Exp Med 194, 991–1002.

Javierre, B.M., Burren, O.S., Wilder, S.P., Kreuzhuber, R., Hill, S.M., Sewitz, S., Cairns, J., Wingett, S.W., Varnai, C., Thiecke, M.J., et al. (2016). Lineage-Specific Genome Architecture Links Enhancers and Non-coding Disease Variants to Target Gene Promoters. Cell 167, 1369–1384 e1319.

Jundt, F., Schwarzer, R., and Dorken, B. (2008). Notch signaling in leukemias and lymphomas. Curr Mol Med 8, 51–59.

Kawamata, S., Du, C., Li, K., and Lavau, C. (2002). Overexpression of the Notch target genes Hes in vivo induces lymphoid and myeloid alterations. Oncogene 21, 3855–3863.

Li, L., Leid, M., and Rothenberg, E.V. (2010a). An early T cell lineage commitment checkpoint dependent on the transcription factor Bcl11b. Science 329, 89–93.

Li, P., Burke, S., Wang, J., Chen, X., Ortiz, M., Lee, S.C., Lu, D., Campos, L., Goulding, D., Ng, B.L., et al. (2010b). Reprogramming of T cells to natural killer-like cells upon Bcl11b deletion. Science 329, 85–89.

Liefke, R., Oswald, F., Alvarado, C., Ferres-Marco, D., Mittler, G., Rodriguez, P., Dominguez, M., and Borggrefe, T. (2010). Histone demethylase KDM5A is an integral part of the core Notch-RBP-J repressor complex. Genes Dev 24, 590–601.

Lindberg, M.J., Popko-Scibor, A.E., Hansson, M.L., and Wallberg, A.E. (2010). SUMO modification regulates the transcriptional activity of MAML1. FASEB J 24, 2396–2404.

Margolin, A.A., Palomero, T., Sumazin, P., Califano, A., Ferrando, A.A., and Stolovitzky, G. (2009). ChIP-on-chip significance analysis reveals large-scale binding and regulation by human transcription factor oncogenes. Proc Natl Acad Sci U S A 106, 244–249.

McManus, S., Ebert, A., Salvagiotto, G., Medvedovic, J., Sun, Q., Tamir, I., Jaritz, M., Tagoh, H., and Busslinger, M. (2011). The transcription factor Pax5 regulates its target genes by recruiting chromatin-modifying proteins in committed B cells. EMBO J 30, 2388–2404.

Medvedovic, J., Ebert, A., Tagoh, H., and Busslinger, M. (2011). Pax5: a master regulator of B cell development and leukemogenesis. Adv Immunol 111, 179–206.

Nutt, S.L., Heavey, B., Rolink, A.G., and Busslinger, M. (1999). Commitment to the B-lymphoid lineage depends on the transcription factor Pax5. Nature 401, 556–562.

Packard, T.A., and Cambier, J.C. (2013). B lymphocyte antigen receptor signaling: initiation, amplification, and regulation. F1000Prime Rep 5, 40.

Palomero, T., Lim, W.K., Odom, D.T., Sulis, M.L., Real, P.J., Margolin, A., Barnes, K.C., O’Neil, J., Neuberg, D., Weng, A.P., et al. (2006). NOTCH1 directly regulates c-MYC and activates a feed-forward-loop transcriptional network promoting leukemic cell growth. Proc Natl Acad Sci U S A 103, 18261–18266.

Popko-Scibor, A.E., Lindberg, M.J., Hansson, M.L., Holmlund, T., and Wallberg, A.E. (2011). Ubiquitination of Notch1 is regulated by MAML1-mediated p300 acetylation of Notch1. Biochem Biophys Res Commun 416, 300–306.

Pridans, C., Holmes, M.L., Polli, M., Wettenhall, J.M., Dakic, A., Corcoran, L.M., Smyth, G.K., and Nutt, S.L. (2008). Identification of Pax5 target genes in early B cell differentiation. J Immunol 180, 1719–1728.

Ryan, R.J.H., Petrovic, J., Rausch, D.M., Zhou, Y., Lareau, C.A., Kluk, M.J., Christie, A.L., Lee, W.Y., Tarjan, D.R., Guo, B., et al. (2017). A B Cell Regulome Links Notch to Downstream Oncogenic Pathways in Small B Cell Lymphomas. Cell Rep 21, 784–797.

Saint Just Ribeiro, M., Hansson, M.L., and Wallberg, A.E. (2007). A proline repeat domain in the Notch co-activator MAML1 is important for the p300-mediated acetylation of MAML1. Biochem J 404, 289–298.

Sanchez-Martin, M., and Ferrando, A. (2017). The NOTCH1-MYC highway toward T-cell acute lymphoblastic leukemia. Blood 129, 1124–1133.

Schmitt, T.M., and Zuniga-Pflucker, J.C. (2002). Induction of T cell development from hematopoietic progenitor cells by delta-like-1 in vitro. Immunity 17, 749–756.

Schwalb, B., Michel, M., Zacher, B., Fruhauf, K., Demel, C., Tresch, A., Gagneur, J., and Cramer, P. (2016). TT-seq maps the human transient transcriptome. Science 352, 1225–1228.

Sharma, V.M., Calvo, J.A., Draheim, K.M., Cunningham, L.A., Hermance, N., Beverly, L., Krishnamoorthy, V., Bhasin, M., Capobianco, A.J., and Kelliher, M.A. (2006). Notch1 contributes to mouse T-cell leukemia by directly inducing the expression of c-myc. Mol Cell Biol 26, 8022– 8031.

South, A.P., Cho, R.J., and Aster, J.C. (2012). The double-edged sword of Notch signaling in cancer. Semin Cell Dev Biol 23, 458–464.

Urbanek, P., Wang, Z.Q., Fetka, I., Wagner, E.F., and Busslinger, M. (1994). Complete block of early B cell differentiation and altered patterning of the posterior midbrain in mice lacking Pax5/BSAP. Cell 79, 901–912.

Van Nguyen, T., Angkasekwinai, P., Dou, H., Lin, F.M., Lu, L.S., Cheng, J., Chin, Y.E., Dong, C., and Yeh, E.T. (2012). SUMO-specific protease 1 is critical for early lymphoid development through regulation of STAT5 activation. Mol Cell 45, 210–221.

Wang, H., Zou, J., Zhao, B., Johannsen, E., Ashworth, T., Wong, H., Pear, W.S., Schug, J., Blacklow, S.C., Arnett, K.L., et al. (2011). Genome-wide analysis reveals conserved and divergent features of Notch1/RBPJ binding in human and murine T-lymphoblastic leukemia cells. Proc Natl Acad Sci U S A 108, 14908–14913.

Wang, Z., Zang, C., Rosenfeld, J.A., Schones, D.E., Barski, A., Cuddapah, S., Cui, K., Roh, T.Y., Peng, W., Zhang, M.Q., et al. (2008). Combinatorial patterns of histone acetylations and methylations in the human genome. Nat Genet 40, 897–903.

Weng, A.P., Ferrando, A.A., Lee, W., Morris, J.P.t., Silverman, L.B., Sanchez-Irizarry, C., Blacklow, S.C., Look, A.T., and Aster, J.C. (2004). Activating mutations of NOTCH1 in human T cell acute lymphoblastic leukemia. Science 306, 269–271.

Weng, A.P., Millholland, J.M., Yashiro-Ohtani, Y., Arcangeli, M.L., Lau, A., Wai, C., Del Bianco, C., Rodriguez, C.G., Sai, H., Tobias, J., et al. (2006). c-Myc is an important direct target of Notch1 in T-cell acute lymphoblastic leukemia/lymphoma. Genes Dev 20, 2096–2109.

Williams, C.J., Naito, T., Arco, P.G., Seavitt, J.R., Cashman, S.M., De Souza, B., Qi, X., Keables, P., Von Andrian, U.H., and Georgopoulos, K. (2004). The chromatin remodeler Mi-2beta is required for CD4 expression and T cell development. Immunity 20, 719–733.

Yang, T., Jian, W., Luo, Y., Fu, X., Noguchi, C., Bungert, J., Huang, S., and Qiu, Y. (2012). Acetylation of histone deacetylase 1 regulates NuRD corepressor complex activity. J Biol Chem 287, 40279–40291.

Yashiro-Ohtani, Y., Wang, H., Zang, C., Arnett, K.L., Bailis, W., Ho, Y., Knoechel, B., Lanauze, C., Louis, L., Forsyth, K.S., et al. (2014). Long-range enhancer activity determines Myc sensitivity to Notch inhibitors in T cell leukemia. Proc Natl Acad Sci U S A 111, E4946–4953.

Yatim, A., Benne, C., Sobhian, B., Laurent-Chabalier, S., Deas, O., Judde, J.G., Lelievre, J.D., Levy, Y., and Benkirane, M. (2012). NOTCH1 nuclear interactome reveals key regulators of its transcriptional activity and oncogenic function. Mol Cell 48, 445–458.

## Supplemental References

Dignam, J.D., Lebovitz, R.M., and Roeder, R.G. (1983). Accurate transcription initiation by RNA polymerase II in a soluble extract from isolated mammalian nuclei. Nucleic Acids Res 11, 1475–1489.

Ienasescu, H., Li, K., Andersson, R., Vitezic, M., Rennie, S., Chen, Y., Vitting-Seerup, K., Lagoni, E., Boyd, M., Bornholdt, J., et al. (2016). On-the-fly selection of cell-specific enhancers, genes, miRNAs and proteins across the human body using SlideBase. Database (Oxford) 2016.

Javierre, B.M., Burren, O.S., Wilder, S.P., Kreuzhuber, R., Hill, S.M., Sewitz, S., Cairns, J., Wingett, S.W., Varnai, C., Thiecke, M.J., et al. (2016). Lineage-Specific Genome Architecture Links Enhancers and Non-coding Disease Variants to Target Gene Promoters. Cell 167, 1369– 1384 e1319.

Kourtis, N., Lazaris, C., Hockemeyer, K., Balandran, J.C., Jimenez, A.R., Mullenders, J., Gong, Y., Trimarchi, T., Bhatt, K., Hu, H., et al. (2018). Oncogenic hijacking of the stress response machinery in T cell acute lymphoblastic leukemia. Nat Med 24, 1157–1166.

Kumar, D., Shadrach, J.L., Wagers, A.J., and Lassar, A.B. (2009). Id3 is a direct transcriptional target of Pax7 in quiescent satellite cells. Mol Biol Cell 20, 3170–3177.

Lawrence, M., Huber, W., Pages, H., Aboyoun, P., Carlson, M., Gentleman, R., Morgan, M.T., and Carey, V.J. (2013). Software for computing and annotating genomic ranges. PLoS Comput Biol 9, e1003118.

Li, H., and Durbin, R. (2009). Fast and accurate short read alignment with Burrows-Wheeler transform. Bioinformatics 25, 1754–1760.

Lizio, M., Harshbarger, J., Shimoji, H., Severin, J., Kasukawa, T., Sahin, S., Abugessaisa, I., Fukuda, S., Hori, F., Ishikawa-Kato, S., et al. (2015). Gateways to the FANTOM5 promoter level mammalian expression atlas. Genome Biol 16, 22.

Love, M.I., Huber, W., and Anders, S. (2014). Moderated estimation of fold change and dispersion for RNA-seq data with DESeq2. Genome Biol 15, 550.

Nakatani, Y., and Ogryzko, V. (2003). Immunoaffinity purification of mammalian protein complexes. Methods Enzymol 370, 430–444.

Phanstiel, D.H., Boyle, A.P., Araya, C.L., and Snyder, M.P. (2014). Sushi.R: flexible, quantitative and integrative genomic visualizations for publication-quality multi-panel figures. Bioinformatics 30, 2808–2810.

Ran, F.A., Hsu, P.D., Wright, J., Agarwala, V., Scott, D.A., and Zhang, F. (2013). Genome engineering using the CRISPR-Cas9 system. Nat Protoc 8, 2281–2308.

Sanjana, N.E., Shalem, O., and Zhang, F. (2014). Improved vectors and genome-wide libraries for CRISPR screening. Nat Methods 11, 783–784.

Stadelmayer, B., Micas, G., Gamot, A., Martin, P., Malirat, N., Koval, S., Raffel, R., Sobhian, B., Severac, D., Rialle, S., et al. (2014). Integrator complex regulates NELF-mediated RNA polymerase II pause/release and processivity at coding genes. Nat Commun 5, 5531.

Thorvaldsdottir, H., Robinson, J.T., and Mesirov, J.P. (2013). Integrative Genomics Viewer (IGV): high-performance genomics data visualization and exploration. Brief Bioinform 14, 178–192.

Wang, N.J., Sanborn, Z., Arnett, K.L., Bayston, L.J., Liao, W., Proby, C.M., Leigh, I.M., Collisson, E.A., Gordon, P.B., Jakkula, L., et al. (2011). Loss-of-function mutations in Notch receptors in cutaneous and lung squamous cell carcinoma. Proc Natl Acad Sci U S A 108, 17761–17766.

Wickham, H. (2009). ggplot2: Elegant Graphics for Data Analysis (New York: Springer-Verlag).

Zhang, Y., Liu, T., Meyer, C.A., Eeckhoute, J., Johnson, D.S., Bernstein, B.E., Nusbaum, C., Myers, R.M., Brown, M., Li, W., et al. (2008). Model-based analysis of ChIP-Seq (MACS). Genome Biol 9, R137.

